# Resilience through diversity: Loss of neuronal heterogeneity in epileptogenic human tissue impairs network resilience to sudden changes in synchrony

**DOI:** 10.1101/2021.03.02.433627

**Authors:** Scott Rich, Homeira Moradi Chameh, Jeremie Lefebvre, Taufik A Valiante

## Abstract

A myriad of pathological changes associated with epilepsy can be recast as decreases in cell and circuit heterogeneity. We thus propose recontextualizing epileptogenesis as a process where reduction in cellular heterogeneity in part, renders neural circuits less resilient to seizure. By comparing patch clamp recordings from human layer 5 (L5) cortical pyramidal neurons from epileptogenic and non-epileptogenic tissue, we demonstrate significantly decreased biophysical heterogeneity in seizure generating areas. Implemented computationally, this renders model neural circuits prone to sudden transitions into synchronous states with increased firing activity, paralleling ictogenesis. This computational work also explains the surprising finding of significantly decreased excitability in the population activation functions of neurons from epileptogenic tissue. Finally, mathematical analyses reveal a unique bifurcation structure arising only with low heterogeneity and associated with seizure-like dynamics. Taken together, this work provides experimental, computational, and mathematical support for the theory that ictogenic dynamics accompany a reduction in biophysical heterogeneity.

## Introduction

Epilepsy, the most common serious neurological disorder in the world (Reynolds, 2002), is characterized by the brain’s proclivity for seizures, which exhibit highly correlated electrophysiological activity and elevated neuronal spiking (Jiruska et al., 2013). While the etiologies that predispose the brain to epilepsy are myriad (Jasper, 2012), the dynamics appear to be relatively conserved (Jirsa et al., 2014; Saggio et al., 2020), suggesting a small palette of candidate routes to the seizure state. One potential route to ictogenesis is disruption of excitatory/inhibitory balance (EIB) - a possible “final common pathway” for various epileptogenic etiologies motivating decades of research into epilepto- and ictogenesis (Dehghani et al., 2016; Žiburkus et al., 2013). A disrupted EIB can impair the resilience of neural circuits to correlated inputs (Renart et al., 2010), a paramount characteristic of ictogenesis. In addition to EIB, biophysical heterogeneity also provides resilience to correlated inputs (Mishra & Narayanan, 2019). Thus, EIB can be considered a synaptic mechanism for input decorrelation, while biophysical heterogeneity contributes to decorrelation post-synaptically.

Cellular heterogeneity is the norm in biological systems (Altschuler & Wu, 2010; Marder & Goaillard, 2006). In the brain, experimental and theoretical work has demonstrated that such heterogeneity expands the informational content of neural circuits, in part by reducing correlated neuronal activity (Padmanabhan & Urban, 2010; Tripathy et al., 2013). Since heightened levels of firing and firing rate correlations hallmark seizures (Jirsa et al., 2014; Zhang et al., 2011), we hypothesize that epilepsy may be likened, in part, to pathological reductions in biological heterogeneity which impair decorrelation, and thus circuit resilience to information poor (Trevelyan et al., 2013), high-firing (Jiruska et al., 2013), and highly-correlated states (Zhang et al., 2011).

A number of pathological changes accompanying epileptogenesis can be recast as decreases in biological heterogeneity. Losses of specific cell-types homogenize neural populations (Cossart et al., 2001; Cobos et al., 2005), down- or upregulation of ion channels homogenize biophysical properties (Arnold et al., 2019; Klaassen et al., 2006; Albertson et al., 2011), and synaptic sprouting homogenizes neural inputs (Sutula & Dudek, 2007). This recontextualizes epileptogenesis as a process associated in part with the progressive loss of biophysical heterogeneity.

To explore this hypothesis we combine electrophysiological recordings from human cortical tissue, computational modeling, and mathematical analysis to detail the existence and consequences of one reduction in biological heterogeneity in epilepsy: the decrease of intrinsic neuronal heterogeneity. We first provide experimental evidence for decreased biophysical heterogeneity in neurons within brain regions that generate seizures (epileptogenic zone) when compared to non-epileptogenic regions. This data constrains an exploration of the effects of heterogeneity in neural excitability on simulated brain circuits. Using a cortical excitatory-inhibitory (E-I) spiking neural network, we show that networks with neuronal heterogeneity mirroring epileptogenic tissue are more vulnerable to sudden shifts from an asynchronous to a synchronous state with clear parallels to seizure onset. Networks with neuronal heterogeneity mirroring non-epileptogenic tissue are more resilient to such transitions. These differing heterogeneity levels also underlie significant, yet counter-intuitive, differences in neural activation functions (i.e., frequency-current or FI curves) measured inside and outside the epileptogenic zone. Using mean-field analysis, we show that differences in the vulnerability to these sudden transitions and activation functions are both consequences of varying neuronal heterogeneities. Viewed together, our experimental, computational, and mathematical results strongly support the hypothesis that biophysical heterogeneity enhances the dynamical resilience of neural networks while explaining how reduced diversity can predispose circuits to seizure-like dynamics.

## Results

### Intrinsic biophysical heterogeneity is reduced in human epileptogenic cortex

In search of experimental evidence for reduced biophysical heterogeneity in epileptogenic regions, we utilized the rare access to live human cortical tissue obtained during resective surgery. Whole-cell current clamp recordings characterized the passive and active properties of layer 5 (L5) cortical pyramidal cells from these samples, a cell type we have shown to display notable biophysical heterogeneity (Moradi Chameh et al., 2021). Biophysical properties of neurons from epileptogenic frontal lobe cortex were contrasted to frontal lobe neurons of tumor patients, with no previous history of seizures, taken a distance from the tumor. Additionally, we obtained, from patients with mesial temporal sclerosis, recordings from neurons in non-epileptogenic middle temporal gyrus (MTG), which is the overlying cortex routinely removed to approach deep temporal structures. The MTG is a well-characterized part of the human brain, representing a common anatomical region from which non-epileptogenic brain tissue has been studied electrophysiologically and transcriptomically (Hodge et al., 2019; Moradi Chameh et al., 2021; Beaulieu-Laroche et al., 2018; Kalmbach et al., 2021), and thus our primary source of non-epileptogenic neurons. We note that each of these studies classify these neurons as indicative of “seemingly normal” human neurons independent of the patients’ epilepsy or tumor diagnoses (i.e., a best case control given limitations in obtaining human tissue).

While multiple sources of heterogeneity were recorded in a variety of physiological measurements (Supplementary Figure S1), we concentrated on attributes of cellular heterogeneity that demonstrated significant differences between the epileptogenic and non-epileptogenic settings. The first was the distance to threshold (DTT) measured as the difference between the resting membrane potential (RMP) and threshold voltage (see Supplementary Figure S1 for these measures presented individually). DTT displayed reduced variability (smaller coefficient of variation (CV); p=0.04; two sample coefficient of variation test) in neurons from epileptogenic frontal lobe (n=13, CV=20.3%) as compared to non-epileptogenic MTG (n=77, CV=37.1%). A significant difference (smaller CV; p=0.03) was also seen when comparing epileptogenic frontal lobe to non-epileptogenic frontal lobe (n=12, CV=40.8%). Meanwhile, the CVs were not significantly different when comparing non-epileptogenic MTG and non-epileptogenic frontal lobe (p=0.7). These features are more easily appreciated from the Gaussian fits of this data presented in Figure 1**(b)**. These results imply that the decrease in biophysical heterogeneity observed in epileptogenic cortex was not confounded by sampling from the temporal versus frontal lobe.

**Figure 1.**
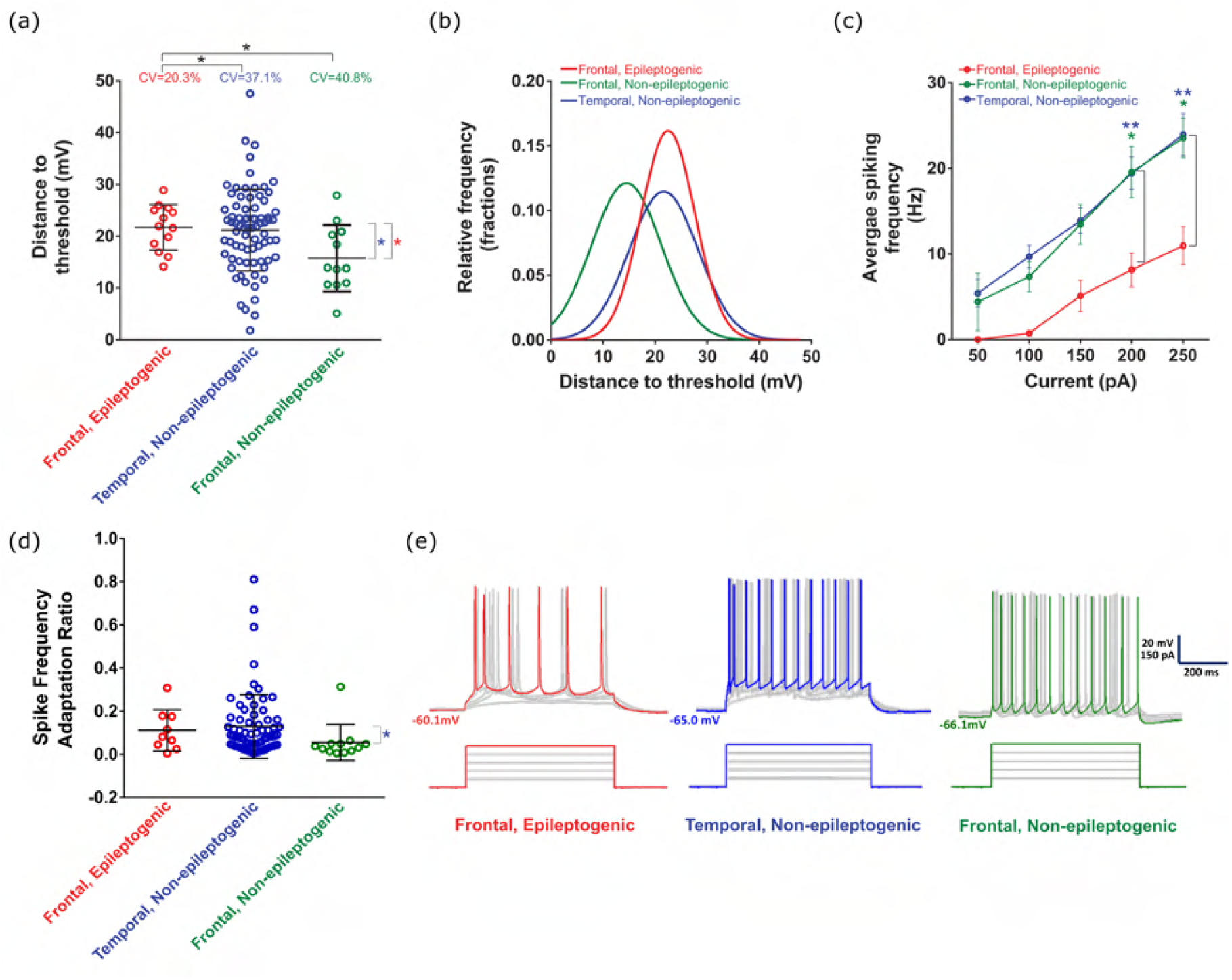
*In vitro* human tissue recordings reveal significantly different electrophysiological heterogeneity between epileptogenic and non-epileptogenic populations. **(a)**: The coefficient of variation (CV) in the distance to threshold (DTT) is significantly larger in both the temporal, non-epileptogenic (i.e., non-epileptogenic MTG; n=77) and frontal, non-epileptogenic (i.e., non-epileptogenic frontal lobe; n=12) populations compared to the frontal, epileptogenic (i.e., epileptogenic frontal lobe; n=13) population (p=0.04 to temporal, non-epileptogenic, p=0.03 to frontal, non-epileptogenic; two sample coefficient of variation test). The CV measure is implemented considering the significantly reduced mean DTT in frontal, non-epileptogenic data compared to the other two populations (p=0.01 for both comparisons; non-parametric Mann-Whitney test). We compare the frontal, epileptogenic and temporal, non-epileptogenic populations computationally given their similar mean DTT (p=0.7). Plotted bars indicate mean *±* standard deviation (SD). **(b)**: An alternative visualization of the DTT distributions via fit Gaussian probability density functions. All three data sets were deemed normal after passing both the Shapiro-Wilk and D’Agostino & Pearson omnibus normality test with alpha=0.05. **(c)**: Neurons from non-epileptogenic populations show similar, linear activation functions (i.e., FI curves). Firing frequency is significantly lower in the frontal, epileptogenic population for a 200 pA injection compared to the temporal, non-epileptogenic (p=0.009; two-way ANOVA-Tukey’s multiple comparison test) and frontal, non-epileptogenic (p=0.03) populations, as well as for a 250 pA injection compared to the temporal, non-epileptogenic (p=0.002) and frontal, non-epileptogenic (p=0.02) populations. Plotted bars indicate mean *±* standard error measure (SEM). **(d)**: All three populations show a similar spike frequency adaptation ratio (see details in Methods), with the only significant difference being between the means from the frontal, non-epileptogenic and temporal, non-epileptogenic populations (p=0.01; One-Way ANOVA post hoc with Dunn’s multiple comparison test). Plotted bars indicate mean *±* SD. **(e)**: Example cell voltage responses following depolarizing current injections (50-250 pA) from all three populations, as used to calculate the FI curve (colors denote population as in previous panels).

While our non-epileptogenic MTG population is larger, this is unavoidable given the availability of human cortical tissue and the additional efforts required to confirm the tissue’s epileptogenic nature (see Discussion). Statistical tests accounting for unequal population sizes were used in comparing the *population CVs and confirmed using the Krishnamoorthy and Lee test, via the R package cvequality* (Marwick & Krishnamoorthy, 2019), that is robust to uneven sample numbers and small sample sizes (Krishnamoorthy & Lee, 2014). Additionally, the significant difference between the standard deviations (SDs) of the DTTs in non-epileptogenic MTG and epileptogenic frontal lobe (p=0.03, Cohen’s d effect size=0.5; F-test; SD=7.8 mV in non-epileptogenic MTG and SD=4.4 mV in epileptogenic frontal lobe) that is implemented in our models has a “moderate” effect size. Finally, we confirmed that the measured heterogeneities are not biased by variability between patients (Supplementary Figure S2), a finding supported by recent multi-patch data in human cortex showing that biophysical properties demonstrate smaller between-subject than within-subject variability (Planert et al., 2021).

The second measure of cellular excitability that demonstrated significant difference between groups was the FI curve (i.e., activation function), which captures the firing rate (F) as a function of input current (I). The FI curve of the population of neurons from the epileptogenic zone displayed qualitative and quantitative differences compared to neurons from both non-epileptogenic MTG and frontal lobe (Figure 1**(c)**). Interestingly, the FI curve shows that pyramidal cells from the epileptogenic zone require more input current to induce repetitive firing, and have overall decreased firing rates for all input currents (p=0.03 when comparing to non-epileptogenic frontal lobe at 200 pA, p=0.02 when comparing to non-epileptogenic frontal lobe at 250 pA, p=0.009 when comparing to non-epileptogenic MTG at 200 pA, and p=0.002 when comparing to non-epileptogenic MTG at 250 pA; two-way ANOVA-Tukey’s multiple comparison test). This non-linear behavior is in strong contrast to the activation functions measured in non-epileptogenic zones, characterized by both higher and more linear changes in firing rates. All three populations show a similar spike frequency adaptation ratio (Figure 1**(d)**), including no significant difference between epileptogenic frontal lobe and non-epileptogenic MTG (the regions focused on in our modeling), indicating that differences in the FI curve are not due to differing adaptation rates. Example firing traces from each population (in response to each of the current steps used in FI curve generation; note that the spike frequency adaptation ratio is calculated from *one* of these steps, chosen as described in the Methods for each individual neuron) are found in Figure 1**(e)**. This increased excitability of the non-epileptogenic populations appears contradictory to the understanding of seizure as a hyperactive brain state, although some prior studies have hinted at this phenomenon (Colder et al., 1996; Schwartzkroin et al., 1983); additionally, the significantly increased first-spike latency in our epileptogenic population (Supplementary Figure S1**(c)**) is further evidence for the decreased single-cell excitability of neurons in this population. We further investigate this in the context of biophysical heterogeneity below.

FI curves from epileptogenic neurons also demonstrated decreased variability: the standard deviations of the frequencies in the epileptogenic population are significantly lower compared to the temporal, non-epileptogenic population at 150 pA (p=0.02, Levene’s test) and at 200 pA (p=0.03), and to the frontal, non-epileptogenic population at 200 pA (p=0.03). Furthermore, the higher input current required to elicit repetitive spiking in our epileptogenic population can be contextualized as a homogenizing feature, as neurons will respond homogeneously (i.e., without spiking) to a larger range of inputs. The smaller slope of the epileptogenic FI curve has a similar effect when repetitive spiking occurs, as changes in the input current will yield smaller changes in the output firing frequency. These findings showcase an additional pattern of decreased heterogeneity in epileptogenic neurons’ spiking behavior.

### Spiking E-I neural networks with epileptogenic levels of excitatory heterogeneity are more vulnerable to sudden changes in synchrony

Given these experimental results, we next computationally explored the effects of the observed differences in biophysical heterogeneity on the transition to a synchronous state akin to the transition to seizure (Zhang et al., 2011). We developed a spiking network model of a cortical microcircuit comprised of recurrently connected excitatory and inhibitory neurons (see details in Methods), motivated in part by the long history of seizure modeling (Kramer et al., 2005; Jirsa et al., 2014) and previous models of decorrelated activity in the cortex (Vogels & Abbott, 2009; Renart et al., 2010; Ostojic, 2014). Our choice of model parameters (see details in Methods) positioned the system near a tipping point at which synchronous activity might arise (Jadi & Sejnowski, 2014a,b; Neske et al., 2015; Rich et al., 2020b) in order to determine the effects of cellular heterogeneity on this potential transition.

We subjected these networks to a slowly linearly increasing external drive to the excitatory cells. This allowed us to observe the dynamics and stability of the asynchronous state, known to be the physiological state of the cortex (Vogels & Abbott, 2009; Renart et al., 2010; Ostojic, 2014), by determining how vulnerable the network is to a bifurcation forcing the system into a state of increased synchrony and firing. A biological analogue for this paradigm would be an examination of whether induced hyper-excitability might drive the onset of seizure-like activity *in vitro*, although such perturbations can more easily be performed continuously (i.e., our linearly increasing external drive) *in silico*.

To facilitate implementing experimentally-derived heterogeneities in our model, we compared epileptogenic frontal lobe with non-epileptogenic MTG given their similar mean DTT values (p=0.7, non-parametric Mann-Whitney test; mean=21.2 mV for non-epileptogenic MTG and mean=21.7 mV for epileptogenic frontal lobe). These populations display significantly different SDs in their DTT values (reported above). Given the definition of our neuron model (rheobases sampled from a normal distribution with with mean 0, see details in Methods), we implement differing heterogeneities by sampling rheobase values for our neural populations from Gaussian distributions with these varying SDs. In this model, the term rheobase refers to the inflection point of the model neuron activation function (see Methods). Heterogeneity in this mathematically-defined rheobase is the *in silico* analogue of heterogeneity in the DTT (i.e., the distribution of rheobases in Figure 2**(c-d)** corresponds to a horizontal shift to a mean of 0 of the DTT distributions in Figure 1**(b)**).

**Figure 2.**
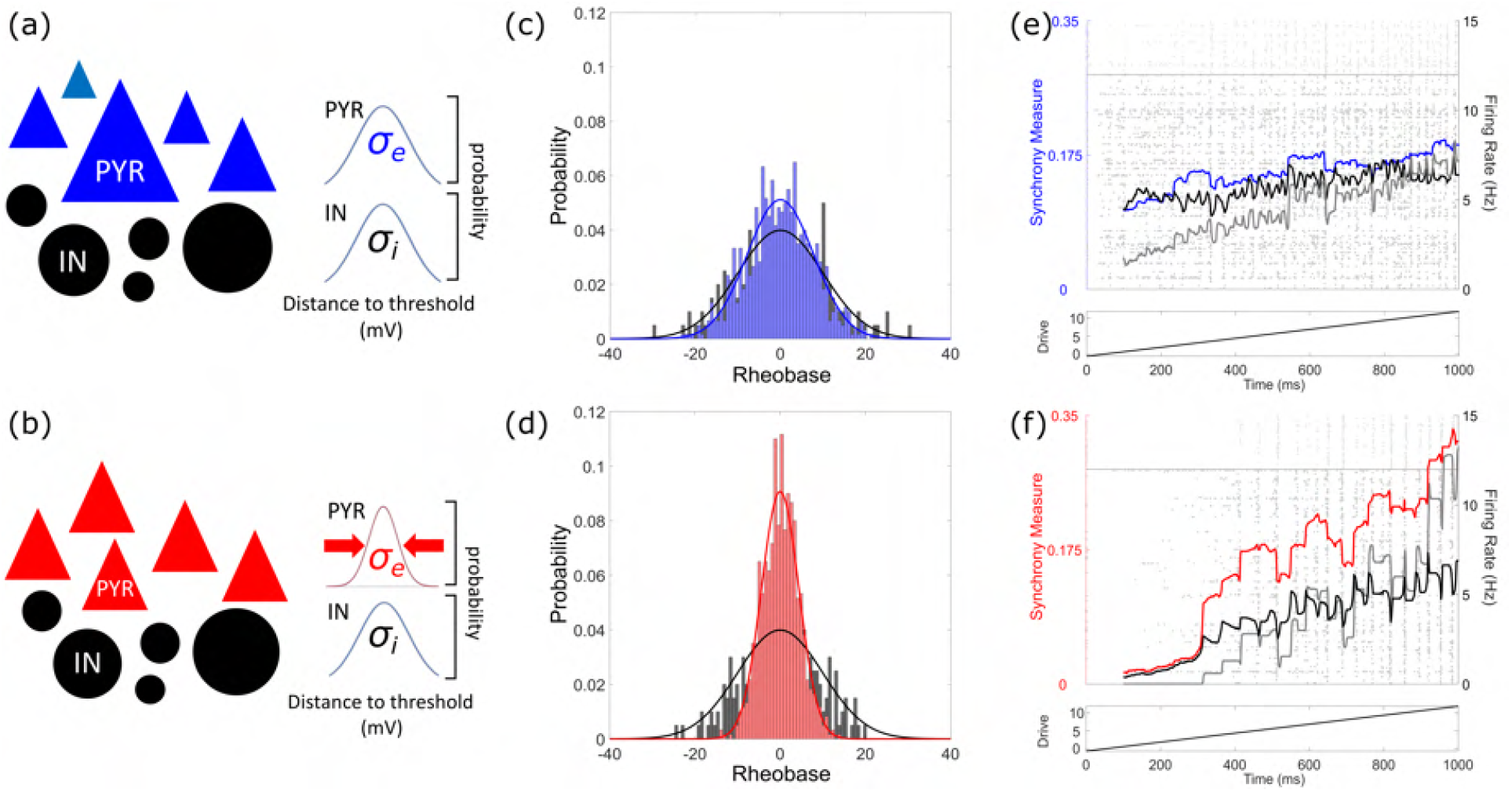
Experimentally observed decreases in heterogeneity amongst excitatory cells promote ictogenic-like transitions in E-I spiking neural network models. **(a-b)**: Schematic representation of model spiking E-I networks, with pyramidal neurons represented as triangles and interneurons as circles. Blue neurons represent non-epileptogenic (i.e. high) levels of heterogeneity (see also the variable neuron sizes) while red neurons represent epileptogenic (i.e. low) levels of heterogeneity (see also the similar neuron sizes). This color schema is maintained in the remaining figures. Here, the inhibitory (black neurons) heterogeneity is set at a moderate value amongst the range studied (*σ*_*i*_ = 10.0 mV), while *σ*_*e*_ = 7.8 mV in panel **(a)** and *σ*_*e*_ = 4.4 mV in panel **(b). (c-d)**: Visualizations of the distribution of model rheobases, with the solid curve (red or blue for excitatory neurons, black for inhibitory neurons) illustrating the Gaussian function and the corresponding histogram illustrating the example random distribution underlying the simulations in this figure. **(e-f)**: Example simulations with a linearly increasing excitatory drive. Background: raster plot of network activity, with each circle representing the firing of an action potential of the associated neuron (excitatory neurons below horizontal line, inhibitory neurons above). Foreground: quantifications of network activity taken over 100 ms sliding time windows, with the excitatory synchrony quantified by the Synchrony Measure in blue or red (left axis), as well as excitatory (black) and inhibitory (grey) population firing rates (right axis). Bottom: drive (*I*(*t*)) to the excitatory population.

The rheobase heterogeneity was parameterized by the SD *σ*_*e*_ for excitatory neurons and *σ*_*i*_ for inhibitory neurons (see diagrams in Figure 2**(a-b)**). This resulted in diversity in the neurons’ activation functions and aligned the variability in their excitabilities with that measured experimentally. We refer to such rheobase heterogeneity simply as heterogeneity in the remainder of the text. Models with non-epileptogenic (high *σ*_*e*_ = 7.8 mV, Figure 2**(e)**) and epileptogenic (low *σ*_*e*_ = 4.4 mV, Figure 2**(f)**) excitatory heterogeneity with identical inhibitory heterogeneity (*σ*_*i*_ = 10.0 mV) exhibit distinct behaviors. With low excitatory heterogeneity, a sharp increase in excitatory synchrony associated with increased firing rates is observed. In contrast, when the excitatory heterogeneity was high, both synchrony and firing rates scaled linearly with input amplitude.

We further investigated the respective roles of excitatory versus inhibitory heterogeneity in these sudden transitions. With non-epileptogenic excitatory heterogeneity (high *σ*_*e*_), increases in excitatory synchrony, excitatory firing rates, and inhibitory firing rates were all largely linear regardless of whether *σ*_*i*_ was low (Figure 3**(a)**) or high (Figure 3**(b)**). Conversely, with excitatory heterogeneity reflective of epileptogenic cortex (low *σ*_*e*_), synchronous transitions were observed for both low (Figure 3**(c)**) and high (Figure 3**(d)**) levels of *σ*_*i*_. This transition is of notably higher amplitude when *σ*_*i*_ is low, indicative of differing underlying dynamical structures driven by *σ*_*i*_.

**Figure 3.**
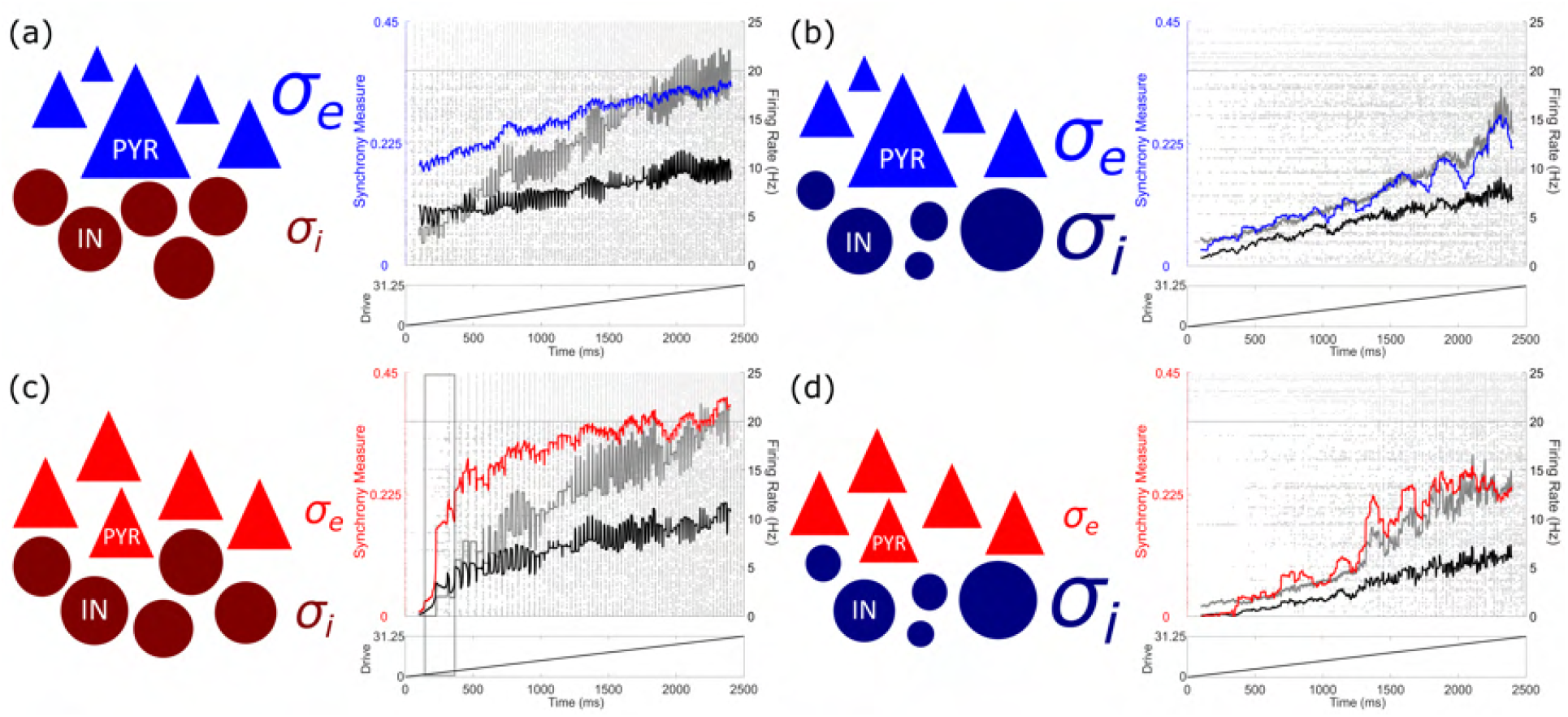
Effects of varied inhibitory heterogeneity on sudden transitions into synchrony in E-I spiking neural network models. Schematics and single simulation visualizations following the conventions of Figure 2 (with inhibitory heterogeneity reflected by darker shaded blue and red neurons), now shown for four combinations of excitatory and inhibitory heterogeneities: *σ*_*e*_ = 7.8 mV and *σ*_*i*_ = 2.5 mV in panel **(a)**, *σ*_*e*_ = 7.8 mV and *σ*_*i*_ = 16.75 mV in panel **(b)**, *σ*_*e*_ = 4.4 mV and *σ*_*i*_ = 2.5 mV in panel **(c)**, and *σ*_*e*_ = 4.4 mV and *σ*_*i*_ = 16.75 mV in panel **(d)**. Relative sizes of *σ*_*e*_ and *σ*_*i*_ represent the relative heterogeneity levels. Transitions into high levels of excitatory synchrony are seen in panel **(c)** and **(d)**, with the transition in panel **(c)** yielding a notably higher level of synchrony (highlighted by the grey box) and occurring much more abruptly. Meanwhile, changes in the dynamics of panels **(a)** and **(b)** are largely linear, with the excitatory synchrony consistently lower when both excitatory and inhibitory heterogeneities are at their highest in panel **(b)**.

Limitations inherent in performing patch-clamp experiments in human cortical tissue prevented the direct measurement of DTT variability in human inhibitory interneurons. To circumvent this, we first studied a range of inhibitory DTT variability aligning with that measured in pyramidal neurons, and then systematically varied and extended this range to account for the possibility of increased heterogeneity amongst the interneuronal population (Cossart, 2011; Huang & Paul, 2019). This enabled the characterization of the contribution of both excitatory and inhibitory heterogeneity to the onset of seizure-like behavior across physiologically relevant ranges of *σ*_*e*_ and *σ*_*i*_. Exploring this range of *σ*_*i*_ values revealed dichotomous dynamics at low and high heterogeneities (Supplementary Figure S3), of which we illustrate exemplars in Figures 3 and 4.

**Figure 4.**
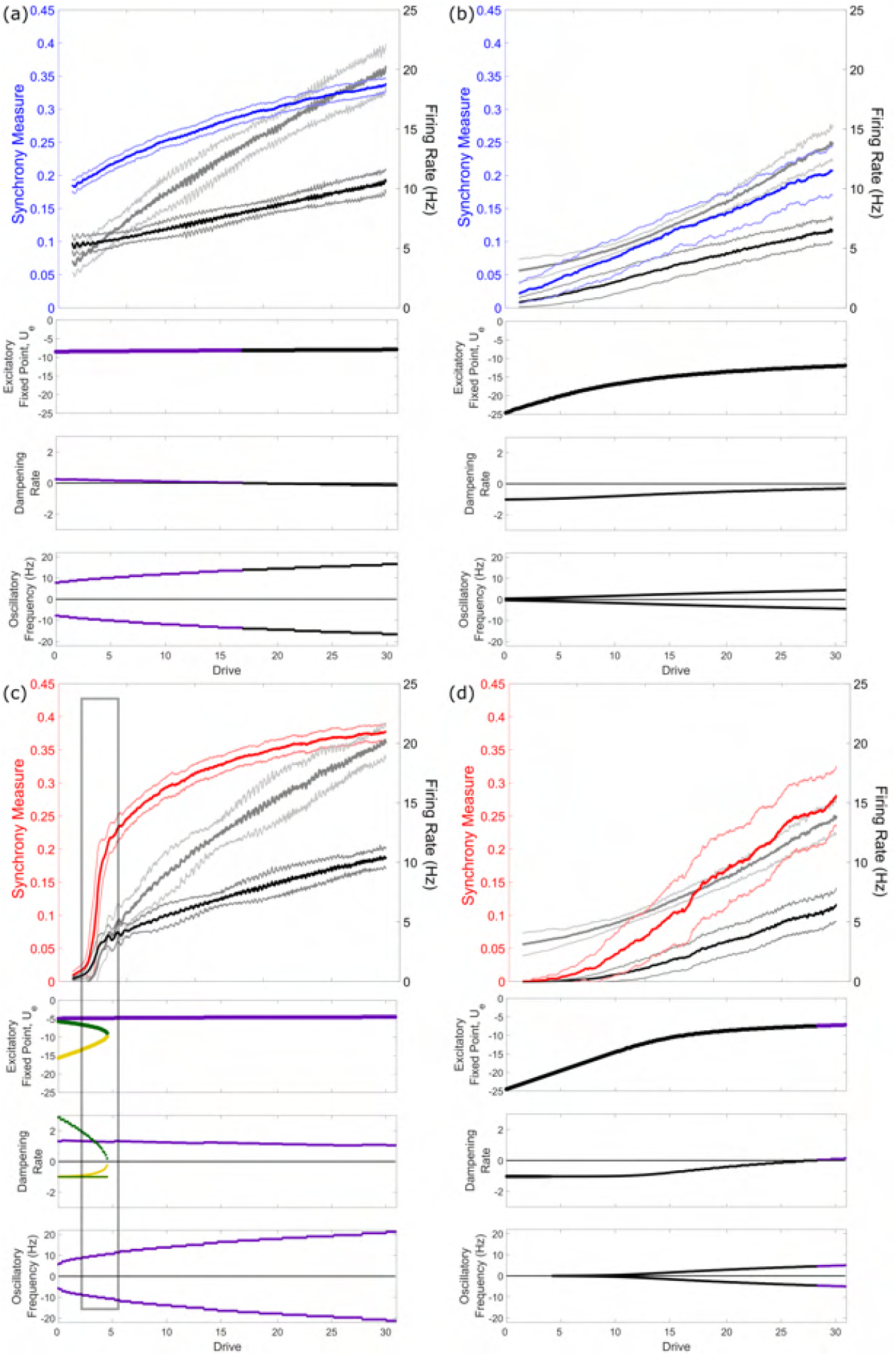
Effects of heterogeneity on spiking network dynamics is explained by stability analysis of mean-field equations. Panels correspond to heterogeneity levels studied in Figure 3. Top row: measures of spiking network dynamics (as seen in Figures 2 and 3) averaged over 100 simulations (dark curve=mean, lighter curve=*±* one SD). Remaining rows: results of stability analysis on mean-field equations corresponding with these networks visualized via the fixed point of mean excitatory activity (top), and the dampening rate and oscillatory frequency associated with each fixed point. Green and gold coloring are used to differentiate the three distinct fixed points in panel c, while the stability of fixed points is color coded (purple=unstable, i.e., positive dampening rate; black=stable, i.e., negative dampening rate). Notably, only in panel **(c)**, where both heterogeneity levels are low, do we see multiple fixed points and a saddle-node bifurcation that occurs at a value of the drive corresponding with the sudden transition in spiking networks (highlighted by the grey box).

### Dynamical differences in networks with varying levels of heterogeneity are explained by their distinct mathematical structures

To gain deeper insight into the effect of heterogeneity at a potential transition to synchrony, we derived and analyzed mathematically the mean-field equations associated with our network model (see Methods). Specifically, we calculated and classified the fixed points of mean-field equations for different values of *σ*_*e*_ and *σ*_*i*_ for the range of drives studied in the spiking networks. The fixed point(s) of the mean-field (for the excitatory population activity, *U*_*e*_) are plotted in the second row of each panel in Figure 4. These values correspond to population averages of the (unitless) membrane potential analogue taken across the individual units in our spiking networks (*u*_*j*_). We then performed linear stability analysis for those fixed points, extracting eigenvalues which determine the fixed points’ stability, and how it might change as input drive is varied. The dampening rate represents the speed at which the system is either repelled from or returns to its fixed point(s) and thus classifies their stability (i.e., the real components of eigenvalues associated with each fixed point). The dampening rate is plotted in the row below the fixed points, followed by the frequency associated with fixed points with imaginary eigenvalues (i.e., the imaginary components of the eigenvalues).

These mean-field analyses confirm that both excitatory and inhibitory heterogeneity have notable impacts on changes in network dynamics analogous to seizure-onset. In the top row of each panel in Figure 4 we present quantifications of our spiking network dynamics as in Figure 3, but averaged over 100 independent simulations. In the presence of high heterogeneity (whenever *σ*_*e*_ and/or *σ*_*i*_ are large, i.e., Figure 4**(a), (b)**, and **(d)**), increased drive results in a smooth and approximately linear increase in both mean activity and synchrony. The mean-field analyses of the associated systems reveal a single fixed point, whose value increases monotonically with drive.

The subtle differences in the spiking network dynamics in these scenarios are reflected in differences in the mean-field analyses. In Figure 4**(d)** a supercritical Hopf bifurcation (Chow & Hale, 2012) at a high level of drive (the stable fixed point becomes unstable, giving rise to a stable limit cycle) is associated with a steeper increase in synchrony. The reverse bifurcation is observed in Figure 4**(a)** (the unstable fixed point becomes stable) and is associated with a slower increase in synchrony, with the synchrony levels being preserved following this bifurcation due to the noise in the spiking networks allowing for the presence of quasi-cycles (Boland et al., 2008). Meanwhile, the fixed point in Figure 4**(b)** is always stable, reflective of the more constant but shallow increase in synchrony in the spiking network.

In contrast to these cases, spiking networks with low heterogeneity (low *σ*_*e*_ and *σ*_*i*_, Figure 4**(c)**) exhibit sudden increases in mean activity and synchrony. The associated mean-field system displays multistability: it possesses multiple fixed points. As the input drive increases, two of these fixed points coalesce and disappear via a saddle-node bifurcation (Chow & Hale, 2012). The system’s mean activity is thus suddenly drawn towards a preexisting large-amplitude limit cycle. This transition occurs at a drive corresponding with the sudden increase in synchrony and mean activity seen in the spiking network. In the mean-field system, the frequency of resulting oscillations are faster compared to the high heterogeneity scenarios, further emphasizing the uniqueness of the dynamical system with low heterogeneity.

We note that the more notable inter-trial variability in Figure 4**(d)** (as illustrated by the fainter ± SD curves) results from the variable (yet gradual) onset of increased synchrony, in contrast to the transition in Figure 4**(c)** which reliably occurs at a specific drive. The different timings of the onset of synchrony in each independent simulation yield oscillations at different relative phases, which explains why oscillations are not observed in our averaged firing rate measures displayed in Figure 4 (notably, such oscillations are subtle even in the single simulation visualizations of Figure 3 given the 100 ms sliding time window); rather, the presence of oscillatory activity is demarcated by a notable increase in the mean Synchrony Measure.

In our mathematical analyses, we focus on characterizing the system’s fixed points and inferring from them the presence of oscillatory behavior associated with limit cycles. Directly identifying such limit cycles is a mathematically arduous process (Savov & Todorov, 2000) unnecessary for the conclusions drawn from our analyses. However, considering the behavior of our spiking networks remains “bounded” (see Supplementary Figure S3**(b)**), we can confidently infer that such limit cycles exist, as is typical when a supercritical Hopf bifurcation yields an unstable fixed point.

To facilitate the comparison of our spiking networks with our mean-field calculations, we developed a Bifurcation Measure (see Methods) quantifying the tendency for sudden (but persistent) changes in the activity of the spiking network. Higher values of this measure indicate the presence of a more abrupt increase in the quantification of interest as the drive increases. Given the more subtle qualitative difference in the firing rates in our spiking networks, we applied the Bifurcation Measure to the excitatory firing rate (*B*_*e*_) for the four combinations of *σ*_*e*_ and *σ*_*i*_ examined in Figure 4. This revealed more sudden changes with low *σ*_*e*_ and *σ*_*i*_ (*B*_*e*_=0.1050) as opposed to any other scenario (high *σ*_*e*_, low *σ*_*i*_, *B*_*e*_=0.0416; high *σ*_*e*_, high *σ*_*i*_, *B*_*e*_=0.0148; low *σ*_*e*_, high *σ*_*i*_, *B*_*e*_=0.0333) where the transition is smoother. This analysis indicates that the dynamical transition present in Figure 4**(c)** is not only unique in the magnitude of the synchronous onset, but also in an associated sudden increase in firing rates.

Since the seizure state is typified both by increased synchrony and firing rates (Jiruska et al., 2013; Zhang et al., 2011), this analysis confirms that the sharp transition in these quantities only observed in spiking models with low heterogeneity is driven by a saddle-node bifurcation (Figure 4**(c)**). These results echo other seizure modeling studies showcasing that ictogenic transitions can arise driven by mathematical bifurcations, and specifically the observation that saddle-node bifurcations underlie abrupt seizure-onset dynamics (Kramer et al., 2005; Jirsa et al., 2014; Saggio et al., 2020). As a corollary, high heterogeneity improves network resilience to sudden changes in synchrony by preventing multistability and fostering gradual changes in network firing rate and oscillatory behavior.

### Asymmetric effects of excitatory and inhibitory heterogeneity

Figure 4 highlights distinct effects of excitatory versus inhibitory heterogeneity on the onset of synchrony in spiking networks and the structure of mean-field systems (see the differences between Figure 4**(a)** and **(c)**). To clarify these effects we explored a larger parameter space of *σ*_*e*_ and *σ*_*i*_, as shown in Supplementary Figure S3. For each heterogeneity combination we applied the Bifurcation Measure to excitatory synchrony (*B*, hereafter referred to simply as the Bifurcation Measure; see details in Methods), which quantifies the abruptness of increased network synchrony in response to a changing network drive. This exploration confirms the asymmetric effect of excitatory and inhibitory heterogeneity on these sudden transitions, with a moderate value of *B* for low *σ*_*e*_ and high *σ*_*i*_ but a minimal value of *B* for high *σ*_*e*_ and low *σ*_*i*_, comporting with patterns observed in previous computational literature (Mejias & Longtin, 2014).

Similar asymmetry is seen in our spiking network dynamics (*B* in Supplementary Figure S3**(a)** and the Synchrony Measure *S* in Supplementary Figure S3**(b)**) and our mean-field systems (the bolded regimes of networks exhibiting multi-stability in Supplementary Figure S3**(a)** and networks exhibiting an unstable fixed point in Supplementary Figure S3**(b)**). We show an example visualization of the fixed points and their classifications in Supplementary Figure S4. Supplementary Figure S5 shows the details of the determination of fixed point stability in Supplementary Figure S3**(b)**.

We further used the Bifurcation Measure to test whether the asymmetric effects of excitatory and inhibitory heterogeneity are generalizable and confirm our system’s robustness. In Supplementary Figure S6 we show the pattern followed by *B* is robust to changes in connectivity density. In the four exemplar cases highlighted in Figures 3 and 4 the dynamics are robust for reasonable changes to the primary parameters dictating our network topology, as shown in Supplementary Figure S7, and similar robustness in the bifurcation structure of the associated mean-field systems is shown in Supplementary Figure S8.

This analysis shows that notable decreases in *B* occur at higher values of *σ*_*i*_ than they do for *σ*_*e*_, a result which has important implications for our understanding of the potentially differing roles of excitatory and inhibitory heterogeneity in seizure resilience. When the loss of specific interneuron types in some epilepsies (Cossart et al., 2001; Cobos et al., 2005) and increases in inhibition (Klaassen et al., 2006) are viewed as homogenizing changes, these computational predictions may help reconcile how both increases and decreases in inhibition may be destablizing to neuronal circuits.

### Differences in population averaged activation functions explained by differences in neuronal heterogeneity

Finally, we return to the counter-intuitive differences in activation functions measured experimentally. As noted previously, the population of neurons from epileptogenic tissue exhibited qualitatively and quantitatively different activation functions via non-linear and hypo-active firing responses (Figure 1**(c)**).

To understand if heterogeneity accounts for these observations, we computed analytically the averaged activation functions of the excitatory populations in our model networks. In Figure 5**(a)**, the experimentally derived firing frequencies from epileptogenic frontal lobe and non-epileptogenic MTG are plotted alongside activation functions of our model populations. For low heterogeneity, the model population’s activation function captured both the non-linear and low firing rate responses measured experimentally for neurons in the epileptogenic zone. The increased excitability and linearity seen experimentally in non-epileptogenic tissue was captured by the averaged activation function for our more heterogeneous model population. This comparison is appropriate considering the FI curve data from Figure 1**(c)** is averaged over the populations of interest, and is thus analogous to the population activation function of our model neurons.

**Figure 5.**
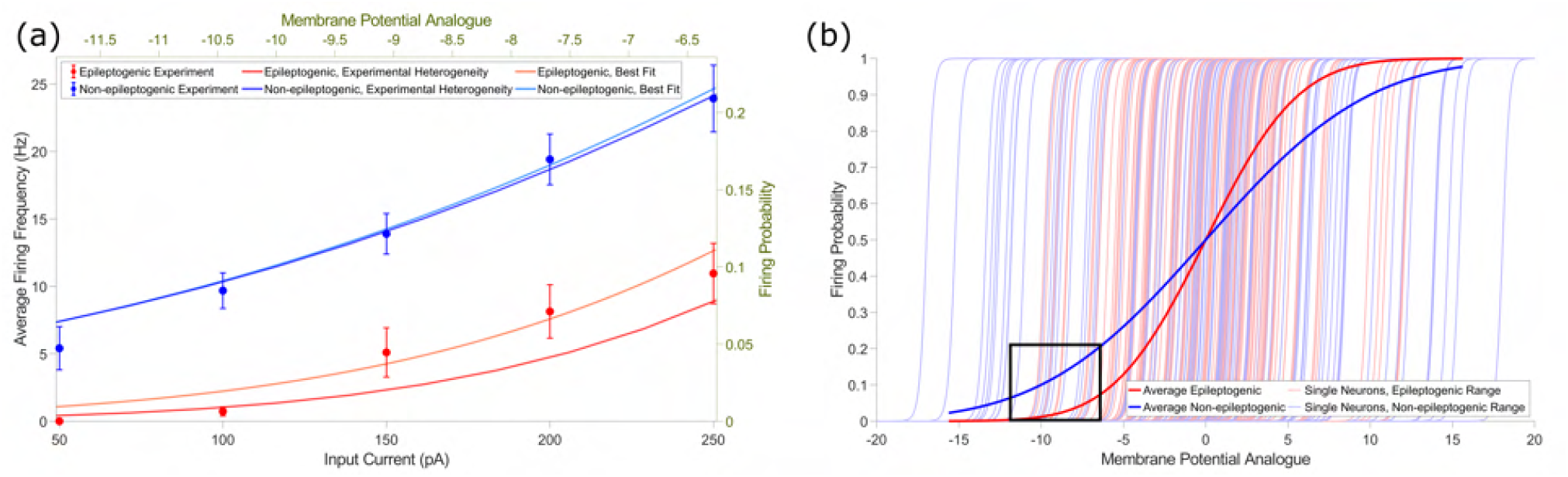
Differing levels of neuronal heterogeneity explain population activation function differences observed experimentally between epileptogenic and non-epileptogenic cortex. **(a)**: Experimentally observed firing frequencies plotted against input current (left and bottom axes, mean *±* SEM) for epileptogenic frontal lobe (red) and non-epileptogenic MTG (blue) tissue (as shown previously in Figure 1**(c)**), visualized against an analogous measure of the relationship between population activity (firing probability) and drive (membrane potential analogue) in our neuron models (right and top axes, details in Methods). The shape of the curve for the heterogeneity value derived from epileptogenic tissue experimentally (red, *σ*_*e*_ = 4.4) qualitatively matches the experimental data, and a best fit (light red, *σ*_*e*_ = 5.03, *r*^2^=0.94) is obtained with a similarly low heterogeneity value. In contrast, the curve associated with the heterogeneity value derived from non-epileptogenic tissue experimentally (blue, *σ*_*e*_ = 7.8) closely matches the experimental data from non-epileptogenic tissue and is nearly identical to the best fit (light blue, *σ*_*e*_ = 7.77, *r*^2^=.98). **(b)**: A visualization of the entirety of the sigmoidal input-output relationship for our neuron models, with the regime compared to experimental data in panel **(a)** in a black box. Fainter curves represent input-output relationships for individual neurons, either epileptogenic (red) or non-epileptogenic (blue): the wider variability in the blue curves yields the flatter sigmoid representing the population activation function for our non-epileptogenic heterogeneity value, and vice-versa for the red curves associated with the epileptogenic heterogeneity value.

To quantitatively support this correspondence, we found the values of *σ*_*e*_ that best fit our experimental data using a non-linear least squares method (see details in Methods). The data from epileptogenic frontal lobe was best fit by an activation function (see Equation 12) with *σ*_*e*_ = 5.0 mV (*r*^2^=0.94), while the data from non-epileptogenic MTG was best fit by an activation function with *σ*_*e*_ = 7.8 mV (*r*^2^=0.98). That the best-fit values closely match the experimentally-observed heterogeneity values means the features of our epileptogenic (resp. non-epileptogenic) activation curves are captured by neural populations with low (resp. high) heterogeneity.

This somewhat counter-intuitive result is explained by the linearizing effect that increased heterogeneity, and noise more generally, has on input-output response functions (Mejias & Longtin, 2014; Lefebvre et al., 2015). This effect is illustrated in Figure 5**(b)**. The bolded sigmoids represent the averaged activity of the entire population of heterogeneous neurons alongside individual activation functions (fainter sigmoids). Increased (resp. decreased) variability dampens (resp. sharpens) the averaged response curve for the non-epileptogenic (resp. epileptogenic) setting. Such variability-induced linearization raises the excitability at low input values, corresponding with the dynamics highlighted in Figure 5**(a)**. Figure 5 illustrates that our model predicts significant differences in the activation function between epileptogenic and non-epileptogenic tissue, and that heterogeneity, or lack thereof, can explain counter-intuitive neuronal responses. However, these differences are not necessarily reflected in network dynamics, as illustrated by the similar network firing rates in Figure 4**(a)** and **(c)** at high levels of drive. In the context of seizure, this implies that excessive synchronization of a neural population need *not* be exclusively associated with increased excitability as represented by a lower minimum input to elicit repetitive firing or higher firing rate of the population of isolated neurons.

## Discussion

In this work, we propose that neuronal heterogeneity may serve an important role in generating resilience to ictogenesis. We explored this hypothesis using *in vitro* electrophysiological characterization of human cortical tissue from epileptogenic and non-epileptogenic areas, which revealed significant differences in DTT (a key determinant of neuronal excitability) variability in the pathological and non-pathological settings. The ability to perform experiments on tissue from human subjects diagnosed with epilepsy makes these results particularly relevant to the human condition. We then implemented these experimentally observed heterogeneities in *in silico* spiking neural networks. Our explorations show that networks with high heterogeneity, similar to the physiological setting, exhibit a more stable asynchronously firing state that is resilient to sudden transitions into a more active and synchronous state. Differing heterogeneity levels also explained the significant differences in the experimentally-obtained population activation functions between epileptogenic and non-epileptogenic tissue. Finally, using mathematical analysis we show that differences in the bifurcation structure of analogous mean-field systems provide a theoretical explanation for dynamical differences in spiking networks. Viewed jointly, these three avenues of investigation provide strong evidence that reduction in biophysical heterogeneity *exists* in epileptogenic tissue, can *yield dynamical changes* with parallels to seizure onset, and that there are *theoretical principles* underlying these differences.

Computational studies have established the role played by heterogeneity in reducing synchronous activity in the context of physiological gamma rhythms (Börgers & Kopell, 2003, 2005; Börgers et al., 2012). Other investigations have implemented heterogeneity in more varied neural parameters (Yim et al., 2013) and identified asymmetric effects of excitatory and inhibitory heterogeneities on network dynamics (Mejias & Longtin, 2012, 2014). Our study complements and extends the understanding of the role of biophysical heterogeneity in neural networks to human epilepsy by: 1) using experimentally derived heterogeneities of the DTT in non-epileptogenic and epileptogenic surgical specimens, which when implemented *in silico* are dynamically relevant; 2) exploring the effects of heterogeneity on the transition to synchrony, a hallmark of seizure onset; 3) detailing the differing extents to which inhibitory and excitatory heterogeneity contribute to circuit resilience to synchronous transitions. Our mathematical analysis further builds on this work to provide a theoretical undergird for these observed dynamics.

The asymmetric effect of excitatory and inhibitory heterogeneities in our model network supports predictions regarding inhibitory heterogeneity’s role in ictogenesis. Supplementary Figure S3**(a)** shows that the sudden onset of synchrony is more likely to arise for moderate values of *σ*_*i*_ than *σ*_*e*_. The physiological heterogeneity of the entire inhibitory population is likely to be larger than for the excitatory population (Cossart, 2011), driven in part by the diverse subpopulations of interneurons (Huang & Paul, 2019). Thus, our work makes two interesting predictions: first, a moderate loss of heterogeneity amongst inhibitory interneurons might be sufficient to make a system vulnerable to ictogenesis; second, the preservation of inhibitory heterogeneity may provide a bulwark against ictogenesis even if excitatory heterogeneity is pathologically reduced as observed experimentally.

Our modeling suggests that post-synaptic inhibitory heterogeneities, in addition to synaptic mechanisms that underlie the decorrelating function of interneurons (Tetzlaff et al., 2012; Sippy & Yuste, 2013), play an important role in the resilience of circuits to sudden transitions to synchronous states. Thus, in addition to changes in EIB (Dehghani et al., 2016; Žiburkus et al., 2013; Jasper, 2012), it is intriguing to speculate that our results might explain both loss (Cobos et al., 2005; Cossart et al., 2001) and gain of function (Klaassen et al., 2006) alterations in inhibition as reduction in interneuronal homogeneity that reduce resilience to ictogenesis.

It is also interesting to conjecture about how these results might be reconciled with the perspective of epilepsy as a disorder of hyper-excitability and the use of high-frequency oscillations (HFOs) as a marker for the epileptogenic zone. Our findings suggest how interictal hypometabolism observed using positron emission tomography (PET) (Niu et al., 2021) and manifestations of “hyper-excitability,” such as inter-ictally recorded HFOs and inter-ictal spikes (IIDs) (Frauscher et al., 2017; Jiruska et al., 2017; Zhang et al., 2011; Schevon et al., 2019), may coexist. We propose that the PET hypometabolism may arise in part from cellular homogenization that reduces population excitability (Figures 1**(c)** and 5**(b)**), since metabolism is tightly linked to firing rate, while this homogenization simultaneously makes the system more vulnerable to transitions into synchronous states (Figure 4**(c)**) such as HFOs, IIDs and seizures.

Notably, previous work has indicated that HFOs arise, in part, from “uninhibited pyramidal cells” (Gulyás & Freund, 2015). Speculatively, this decreased inhibition could arise from a homogenized, and in turn hypo-excitable, inhibitory population (Figure 5). This may further explain the hypometabolism observed inter-ictally given that interneuronal spiking appears to contribute more to brain metabolism than pyramidal cells (Ackermann et al., 1984). While speculative, the interconnected nature of neural heterogeneity and excitability identified in this work can, at minimum, motivate further studies using targeted patching of interneurons in both human and chronic rodent models to characterize if homogenization occurs in interneuronal populations during epileptogenesis and epilepsy.

Our results include fewer neurons from the frontal lobe considering it is a less common source of human cortical tissue than non-epileptogenic MTG. Thus, we use the population of non-epileptogenic frontal lobe neurons only as evidence that heterogeneity levels are not confounded by comparison between the temporal and frontal lobes. The sample size of our epileptogenic neurons was limited by the necessity to confirm the epileptogenicity of the resected cortex using using electrocorticography (ECoG), making this data set highly selective. Although one might obtain a greater sample by comparing non-epileptogenic MTG to epileptogenic mesial temporal structures (i.e., subiculum, parahippocampal gyrus, hippocampus) comparing the allocortex and neocortex would add a further confound. Alternatively, obtaining non-epileptogenic medial temporal lobe (MTL) cortex is exceedingly rare. With these important limitations in the access to human cortical tissue considered, our comparison between epileptogenic frontal lobe, non-epileptogenic frontal lobe, and non-epileptogenic MTG represent a best-case comparison of the biophysical properties of epileptogenic and non-epileptogenic human tissue while controlling for confounds introduced by the differing brain regions. Our computational and mathematical explorations optimize the conclusions that can be drawn from this rare data.

Our model networks, while analogous to E-I microcircuits commonly used in computational investigations of cortical activity (Renart et al., 2010; Ostojic, 2014; Vogels & Abbott, 2009), are simplified from the biophysical reality and are correspondingly limited. For instance, such models cannot reasonably capture the full richness and complexity of seizure dynamics and do not include multiple inhibitory populations (Huang & Paul, 2019). However, this simplifying choice facilitates findings that have their foundation in fundamental mathematical principles and are not especially reliant on biophysical intricacies such as network topology (see the confirmation of the robustness of our models in Supplementary Figures S7 and S8). In addition, experimental limitations arising from patch-clamp experiments limit the number of potential interneurons that can be patched in human tissue, precluding measuring inhibitory DTT and its variability experimentally. Thus, the values of *σ*_*i*_ studied in our model networks were chosen to approximately align with that seen experimentally in the excitatory population while accounting for the possibility of increased inhibitory heterogeneity (Cossart, 2011; Huang & Paul, 2019), with this parameter systematically varied throughout the study.

These limitations warrant the development of biophysically detailed, human inspired neuron and network models, allowing for the study of additional types of heterogeneity. Such studies will benefit from our recent development of a biophysically-detailed computational model of a human L5 cortical pyramidal neuron (Rich et al., 2021). In this vein, while we do not model seizures per se in this work, the two most common types of seizure onsets observed in intracranial recordings are the low-voltage fast (Lee et al., 2000) and hyper-synchronous onsets (Velascol et al., 1999). Both reflect a sudden transition from a desynchronized state to a synchronous oscillation, albeit of differing frequencies. Given the ubiquity of such onsets, our modeling of the transition to synchrony is likely to be broadly relevant to epilepsy.

Lastly, one might wonder what neurobiological processes render an epileptogenic neuronal population less biophysically diverse. While under physiological conditions channel densities are regulated within neurons to obtain target electrical behaviors (Marder, 2011), it remains speculative as to what processes might lead to pathological homogenization of neuronal populations. However, modeling suggests that biological diversity may be a function of input diversity, and thus “homogenizing the input received by a population of neurons should lead the population to be less diverse” (Tripathy et al., 2013), possibly through intrinsic plasticity mechanisms (Beck & Yaari, 2008; Zhang & Linden, 2003). Although requiring further exploration, it is possible that the information-poor, synchronous post-synaptic barrages accompanying a seizure (Trevelyan et al., 2013) represent such a homogenized input, reducing a circuit’s resilience to synchronous transitions and promoting epileptogenesis by reducing biophysical heterogeneity.

## Acknowledgments

We thank Frances Skinner, Shreejoy Tripathy, Prajay Shah, and Anukrati Nigam for productive intellectual discussions on this topic in the project’s early stages. We thank the National Sciences and Engineering Research Council of Canada (NSERC Grants RGPIN-2017-06662 to J.L. and RGPIN-2015-05936 to T.A.V.), the Krembil Foundation (Krembil Seed Grant to J.L. and T.A.V.), the University of Toronto Department of Physiology (Yuet Ngor Wong Award to S.R.), and the Savoy Foundation (Steriade-Savoy Postdoctoral Fellowship to S.R.) for support of this research.

## Author Contributions

Conception and design: SR, HMC, JL, TAV. Experimental data collection: HMC. Data analysis and interpretation: SR, HMC, TAV. Simulations: SR. Mathematical analysis: SR, JL. Initial drafting: SR. Edits and revisions: SR, HMC, JL, TAV. All authors approved the version to be submitted.

## Materials and Methods

### Experiment: Human brain slice preparation

All procedures on human tissue were performed in accordance with the Declaration of Helsinki and approved by the University Health Network Research Ethics board. Patients underwent a standardized temporal or frontal lobectomy under general anesthesia using volatile anesthetics for seizure treatment (Valiante, 2009). Tissue was obtained from patients diagnosed with temporal or frontal lobe epilepsy who provided written consent. Tissue from temporal lobe was obtained from 22 patients, age ranging between 21 to 63 years (mean age ± SEM: 37.8 ± 2.9), with 1-9 cells studied per patient. The resected temporal lobe tissue displayed no structural or functional abnormalities in preoperative MRI and was deemed “healthy” tissue considering it is located outside of the epileptogenic zone. Tissue from epileptogenic frontal lobe was obtained from five patients, age ranging between 23-36 years (mean age ± SEM: 30.2 ± 2.4), and was deemed “epileptogenic” tissue as confirmed using electrocorticography (ECoG), making this data set highly selective. 1-5 cells were studied per patient. Tissue from non-epileptogenic frontal lobe obtained during tumor resection was obtained from two patients, ages 37 and 58 years, with 8 and 4 cells studied per patient, and was also considered “healthy, non-epileptogenic” tissue as it was taken away from the tumor itself. This tissue is a common source of human cortical tissue to study human cell and circuit properties (Kalmbach et al., 2018, 2021; Testa-Silva et al., 2014).

After surgical resection, the cortical tissue block was instantaneously submerged in ice-cold (∼4^*◦*^C) cutting solution that was continuously bubbled with 95% O_2_-5% CO_2_ containing (in mM): sucrose 248, KCl 2, MgSO_4_.7H_2_O 3, CaCl_2_.2H_2_O 1, NaHCO_3_ 26, NaH_2_PO_4_.H_2_O 1.25, and D-glucose 10. The osmolarity was adjusted to 300-305 mOsm. The human tissue samples were transported (5-10 min) from Toronto Western Hospital (TWH) to the laboratory for further slice processing. Transverse brain slices (400 *µ*m) were obtained using a vibratome (Leica 1200 V) perpendicular to the pial surface to ensure that pyramidal cell dendrites were minimally truncated (Beaulieu-Laroche et al., 2018; Kalmbach et al., 2018) in the same cutting solution as used for transport. The total duration, including slicing and transportation, was kept to a maximum of 20-30 minutes. After sectioning, the slices were incubated for 30 min at 34^*◦*^C in standard artificial cerebrospinal fluid (aCSF) (in mM): NaCl 123, KCl 4, CaCl_2_.2H_2_O 1, MgSO_4_.7H_2_O 1, NaHCO_3_ 26, NaH_2_PO_4_.H_2_O 1.2, and D-glucose 10. The pH was 7.40 and after incubation the slice was held for at least for 60 min at room temperature. aCSF in both incubation and recording chambers were continuously bubbled with carbogen gas (95% O_2_-5% CO_2_) and had an osmolarity of 300-305 mOsm.

### Experiment: Electrophysiological recordings and intrinsic physiology feature analysis

Slices were transferred to a recording chamber mounted on a fixed-stage upright microscope (Axioskop 2 FS MOT; Carl Zeiss, Germany). Recordings were performed from the soma of pyramidal neurons at 32-34^*◦*^ in recording aCSF continually perfused at 4 ml/min. Cortical neurons were visualized using an IR-CCD camera (IR-1000, MTI, USA) with a 40x water immersion objective lens. Using the IR-DIC microscope, the boundary between layer 1 (L1) and 2 (L2) was easily distinguishable in terms of cell density. Below L2, the sparser area of neurons (L3) was followed by a tight band of densely packed layer 4 (L4) neurons, with a decrease in cell density indicating layer 5 (L5) (Moradi Chameh et al., 2021; Kalmbach et al., 2021).

Patch pipettes (3-6 MΩ resistance) were pulled from standard borosilicate glass pipettes (thin-wall borosilicate tubes with filaments, World Precision Instruments, Sarasota, FL, USA) using a vertical puller (PC-10, Narishige). Pipettes were filled with intracellular solution containing (in mM): K-gluconate 135; NaCl 10; HEPES 10; MgCl_2_ 1; Na_2_ATP 2; GTP 0.3, pH adjusted with KOH to 7.4 (290–309 mOsm).

Whole-cell patch-clamp recordings were obtained using a Multiclamp 700A amplifier, Axopatch 200B amplifier, pClamp 9.2 and pClamp 10.6 data acquisition software (Axon instruments, Molecular Devices, USA). Electrical signals were digitized at 20 kHz using a 1320X digitizer. The access resistance was monitored throughout the recording (typically between 8-25 MΩ), and neurons were discarded if the access resistance was *>*25 MΩ. The liquid junction potential was calculated to be -10.8 mV and was not corrected.

Electrophysiological data were analyzed off-line using Clampfit 10.7, Python and MATLAB (MATLAB, 2019). Electrophysiological features were calculated from responses elicitepd by 600 ms square current steps as previously described (Moradi Chameh et al., 2021). Briefly, the resting membrane potential (RMP) was measured after breaking into the cell (IC=0). The firing threshold was determined following depolarizing current injections between 50 to 250 pA with 50 pA step size for 600 ms; the threshold was calculated by finding the voltage value corresponding with a value of 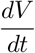 that was 5% of the average maximal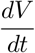 across all action potentials elicited by the input current that first yielded action potential firing. The distance to threshold presented in this paper was calculated as the difference between the RMP and threshold. The average FI curve (i.e., activation function) was generated by calculating the instantaneous frequency at each spike for each of the depolarizing current injections (50-250 pA, step size 50 pA, 600 ms) and averaging over the population. Spike frequency adaptation ratio was calculated from the first current injection that yielded at least four spikes, and is defined as the mean of the ratio of subsequent inter-spike intervals. This could not be quantified in every neuron if sufficient spiking was not elicited by the current-clamp protocol. This analysis utilizes the IPFX package made available through the Allen Institute (https://github.com/AllenInstitute/ipfx), as used by Berg et al. (2021) amongst others.

Plotting of experimental data was performed using GraphPad Prism 6 (GraphPad software, Inc, CA, USA). The non-parametric Mann-Whitney test was used to determine statistical differences between the means of two groups. The F-test was used to compare standard deviation (SD) between groups. The two sample coefficient of variation test was used to compare the coefficient of variance (CV) between groups. Normality of the data was tested with the Shapiro-Wilk and D’Agostino & Pearson omnibus normality tests with alpha=0.05. The one-way ANOVA post hoc with Dunn’s multiple comparison test was used to determine statistical significance in the spike frequency adaptation ratio. A standard threshold of p*<*0.05 is used to report statistically significant differences.

### Modeling: spiking neural network

The cortical spiking neural network contains populations of recurrently connected excitatory and inhibitory neurons (Snyder & Miller, 2012; Stevens & Zador, 1996). The spiking response of those neurons obeys the non-homogeneous Poisson process

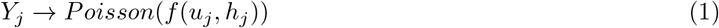

where *Y*_*j*_ = Σ_*l*_ *δ*(*t* − *t*_*k*_) is a Poisson spike train with rate *f* (*u*_*j*_, *h*_*j*_).

The firing rate of neuron *j* is determined by the non-linear sigmoidal activation function *f* (*u*_*j*_, *h*_*j*_),

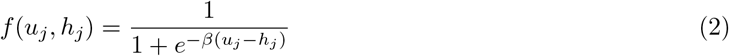

where *u*_*j*_ is the membrane potential analogue and *h*_*j*_ represents the rheobase. The constant *β* = 4.8 scales the non-linear gain.

Heterogeneity is implemented via the rheobases *h*_*j*_. The *h*_*j*_ values are chosen by independently and randomly sampling a normal Gaussian distribution whose standard deviation is *σ*_*e,i*_ if neuron *j* is excitatory (*e*) or inhibitory (*i*). The values of *σ*_*i*_ and *σ*_*e*_ are varied throughout these explorations between a minimum value of 2.5 mV and a maximum value of 16.75 mV. The heterogeneity parameters for the model have a direct parallel with the heterogeneity in the distance to threshold (DTT) measured experimentally, with *β* chosen so that the experimentally observed heterogeneity values and the heterogeneity parameters implemented in the model are within the same range (compare Figure 1**(b)** and Figure 2**(c-d)**).

The membrane potential analogue *u*_*j*_ is defined by

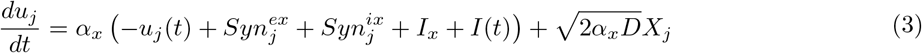

The variable *α*_*x*_ represents the time constant depending upon whether the neuron *j* is excitatory (*x* = *e, α*_*e*_ = 10 ms) or inhibitory (*x* = *i, α*_*i*_ = 5 ms). The differential time scales are implemented given the different membrane time constants between cortical pyramidal neurons and parvalbumin positive (PV) interneurons (Neske et al., 2015).

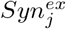 and 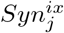 are the synaptic inputs to the cell *j* (from the excitatory and inhibitory populations, respectively), dependent upon whether cell *j* is excitatory (*x* = *e*) or inhibitory (*x* = *i*). Our cortical model is built of 800 excitatory and 200 inhibitory neurons (Traub et al., 1997; Rich et al., 2017, 2018). The connectivity density for each connection type (E-E, E-I, I-E, and I-I) is varied uniformly via a parameter *p*. In this study, *p* = 1 is used (i.e., all-to-all connectivity) with the exception of in Supplementary Figure S6. The synaptic strengths are represented by *w*_*xy*_ where *x, y* = *e, i* depending upon whether the pre-synaptic cell (*x*) and the post-synaptic cell (*y*) are excitatory or inhibitory. In our model, *w*_*ee*_ = 100.000, *w*_*ei*_ = 187.500, *w*_*ie*_ = −293.750, and *w*_*ii*_ = −8.125. Negative signs represent inhibitory signalling, while positive signs represent excitatory signalling. These values are chosen to place the network near a tipping point between asynchronous and synchronous firing based on mathematical analysis and previous modeling work (Rich et al., 2020b), and scaled relative to the values of *β*.

The post-synaptic inputs 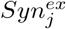 and 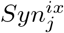 are given by

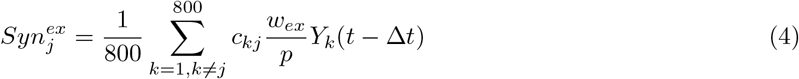

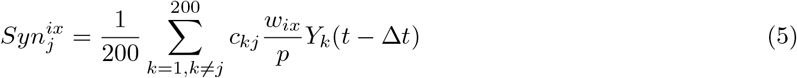

where *x* = *e, i* and *Y*_*k*_ is a Poisson spike train given by *Y*_*k*_ = Σ_*l*_ *δ*(*t* − *t*_*l*_). The connectivity scheme excludes auto-synapses. *c*_*kj*_ represents the connectivity: if neuron *k* synapses onto neuron *j, c*_*kj*_ = 1, and otherwise *c*_*kj*_ = 0. The synaptic weights are scaled by the connectivity density *p* so that the net input signal to each neuron is not affected by the number of connections.

Equation 3 includes three non-synaptic inputs to the neuron: *I*_*x*_, *I*(*t*), and and 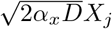. The variable *X*_*j*_ is a spatially independent Gaussian white noise process. The value of noise intensity was chosen so that the noise-induced fluctuations are commensurate with endogenous dynamics of the network. *I*_*x*_ represents a bias current whose value depends on whether the neuron is excitatory (*x* = *e*) or inhibitory (*x* = *i*), imparting a differential baseline spiking rate to these distinct populations. Here, *I*_*i*_ = −31.250, ensuring that inhibitory neurons will typically require excitatory input to fire, matching biophysical intuition. *I*_*e*_ = −15.625 is based on previous literature (Jadi & Sejnowski, 2014a,b; Neske et al., 2015; Rich et al., 2020b) to position the system near the transition between asynchronous and synchronous firing.

*I*(*t*) implements time-varying external input only applied to the excitatory population (this is simply referred to as the “drive” to the system in Figures 2, 3 and 4). In this work, this term is used primarily to study the response of the spiking network to a linear ramp excitatory input that occurs at a time scale much slower than the dynamics of individual neurons: to yield the ramp current used throughout the study *I*(*t*) simply varies linearly between 0 and 31.25 over a 2500 ms simulation (for computational efficiency, the simulation length is limited to 2048 ms for the heatmaps displayed in Supplementary Figures S3 and S6). In Supplementary Figure S3**(b)**, where we characterize the dynamics of the network with constant input, *I*(*t*) = 15.625 uniformly.

The final probability of a Poisson neuron *j* firing at time *t* depends upon the effect of these various elements on *u*_*j*_:

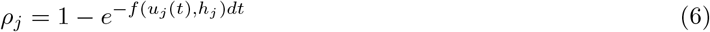

#### Parameter values

Parameter values summarized in Table 1 below are analogous to those used in previous work on oscillatory cortical networks (Jadi & Sejnowski, 2014a,b; Neske et al., 2015; Rich et al., 2020b) with the scaling of our chosen *β* accounted for.

**Table 1.** Key model parameters

| Parameter | Value |
| --- | --- |
| Number of excitatory neurons | 800 |
| Number of inhibitory neurons | 200 |
| Excitatory time constant, $\alpha_e$ | 10 ms |
| Inhibitory time constant, $\alpha_i$ | 5 ms |
| Non-linear gain of activation function, $\beta$ | 4.8 |
| Variance of noisy input, $D$ | 3.906 |
| Excitatory bias current, $I_e$ | -15.625 |
| Inhibitory bias current, $I_i$ | -31.250 |
| External input, $I(t)$ | Variable |
| Excitatory-excitatory synaptic strength, $w_{ee}$ | 100.000 |
| Excitatory-inhibitory synaptic strength, $w_{ei}$ | 187.500 |
| Inhibitory-inhibitory synaptic strength, $w_{ii}$ | -8.125 |
| Inhibitory-excitatory synaptic strength, $w_{ie}$ | -293.750 |
| Excitatory heterogeneity, $\sigma_e$ | Variable |
| Inhibitory heterogeneity, $\sigma_i$ | Variable |
| rheobase, $h$ | Variable |
| Connectivity density, $p$ | Variable |
| Time step, $\Delta t$ | 1 ms |

#### Numerics

All sampling from standard normal Gaussian distributions is done via the Box-Mueller algorithm (Golder & Settle, 1976). Equations are integrated using the Euler-Maruyama method. In our simulations, Δ*t* = 0.1, scaled so that each time step Δ*t* represents 1 ms.

The excitatory network synchrony (i.e. Synchrony Measure) and excitatory and inhibitory firing rates are calculated over sliding 100 ms time windows in Figures 2, 3 and 4. To preserve symmetry and ensure initial transients do not skew the data, our first window begins at *t* = 100.

The Synchrony Measure is an adaptation of a commonly used measure developed by Golomb and Rinzel (Golomb & Rinzel, 1993, 1994) to quantify the degree of coincident spiking in a network as utilized in our previous studies (Rich et al., 2016, 2017, 2018, 2020a). Briefly, the measure involves convolving a very narrow Gaussian function with the time of each action potential for every cell to generate functions *V*_*i*_(*t*). The population averaged voltage *V* (*t*) is then defined as 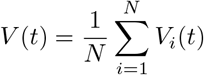, where *N* is the number of cells in the network. The overall variance of the population averaged voltage Var(*V*) and the variance of an individual neuron’s voltage Var(*V*_*i*_) is defined as

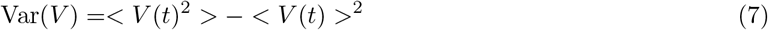

and

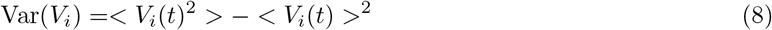

where *<* · *>* indicates time averaging over the interval for which the measure is taken. The Synchrony Measure *S* is then defined as

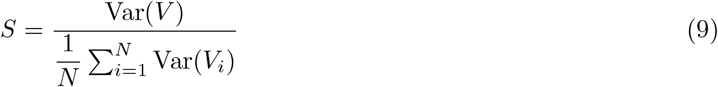

The value *S* = 0 indicates completely asynchronous firing, while *S* = 1 corresponds to fully synchronous network activity. Intermediate values represent intermediate degrees of synchronous firing.

In the case of sliding time bins, this measure is taken by only considering spikes falling into the time window of interest. In Figure 4 we present averages of *S* over 100 independent realizations, and if a particular run yields a “NaN” result for *S* at a given time step (indicating no spikes in the associated window), we eliminate that value from the average for that time point (this increases the variability of these values since there are less to average over; thus, this is reflected in an increased range of the ± STD curves). In contrast, in Supplementary Figure S3**(b)** we generate a single value the Synchrony Measure (or the other measures of interest) over the last 1000 ms of the simulation. Supplementary Figure S3**(b)** displays this measure averaged over five independent simulations.

Supplementary Figure S3 includes the presentation of our Bifurcation Measure *B*. This quantifies the presence of sudden and significant changes in the Synchrony Measure over time. First, we take the Synchrony Measure time series for each independent run (i.e., as presented in Figure 3), and use the *smooth* function in MATLAB(MATLAB, 2019) with a 500 step window, generating a new time series from this moving average filter. This low-pass filter serves to account for fluctuations arising when, for example, a particular 100 ms window includes more or less activity than average. We denote this filtered time-series *S*_*s*_. Second, we calculate the difference quotient 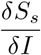, where *I* is the value of the external drive (plotted against time in Figure 3), at each step in the time series. Finally, we take the variance of the values of 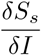 using the *var* function in MATLAB (MATLAB, 2019): networks in which the Synchrony Measure changes in a consistently linear fashion will have a tight distribution of 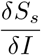 around the average slope (see, for example, Figure 3**(b)**), and thus a low variance; in contrast, networks in which the Synchrony Measure undergoes abrupt transitions will yield a multi-modal distribution of 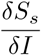, with each mode corresponding to different linear sections of *S*, and thus the variance of these values will be notably higher (see, for example, Figure 3**(c)**). The plotted value of *B* represents an average over the *B* values calculated for each independent network instantiation. We note that when we calculate the “firing rate Bifurcation Measure” *B*_*e*_ in reference to the four scenarios in Figure 4, we simply replicate the above steps on the firing rate time series rather than the Synchrony Measure time series.

We emphasize that the Bifurcation Measure is appropriate for identifying the dynamics of interest in this work given that the related quantifications increase largely monotonically in response to increased drive, especially once these time series are “smoothed” prior to the application of this measure. The smoothed Synchrony Measure and firing rates do not display any discontinuous behaviors in our experimental paradigms that might confound this measure.

#### Analysis of FI curves

In Figure 5, we compare activation functions derived from experimental data with model analogues (i.e., the function *F* described below in Equation 12). In Figure 5**(b)** we show examples of *F* with epileptogenic and non-epileptogenic levels of heterogeneity alongside samples of the function *f* (Equation 2) randomly chosen based on the differing heterogeneity levels.

In Figure 5**(a)**, we confirm the correspondence between the *F* functions and the experimental data by determining the value of *σ*_*e*_ best fitting this data. This process involved three steps: first, we qualitatively determined the portion of the *F* curves most likely to fit this data as that in −11.875 ≤ *U*_*e*_ ≤ −6.25; second, both the x (*U*_*e*_, [-11.875 -6.25]) and y (probability of firing, [0.003585 .2118]) variables were re-scaled to match the ranges exhibited by the x (input current, pA, [50 250]) and y (firing frequency, Hz, [0 24]) variables in the experimental data; finally, a fit was calculated using MATLAB’s (MATLAB, 2019) Curve Fitting application. This process used a non-linear least squares method, with *r*^2^ *>* .93 for both fits (see details in Results). Additional scaling was performed for plotting so that the two x- and y-axes in Figure 5 remain consistent.

### Modeling: Mean-field reduction

Following previous work (Hutt et al., 2016; Stefanescu et al., 2012; Hutt et al., 2020; Rich et al., 2020b; Lefebvre et al., 2015; Hutt et al., 2020) we perform a mean-field reduction of the spiking network in Equation 3. We assume that the firing rate of cells is sufficiently high to make use of the diffusion approximation (Gluss, 1967), yielding

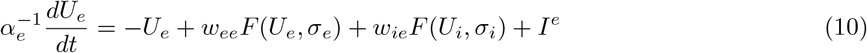

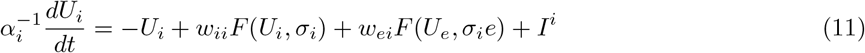

where 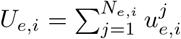 represents the mean activity of the excitatory or inhibitory population, respectively.

The function *F* represents the average activation function conditioned upon the value of *σ*_*e,i*_ via the convolution

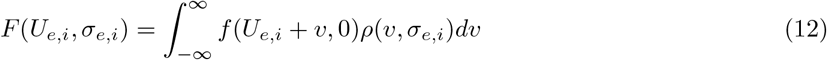

where 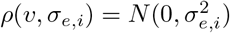 (Lefebvre et al., 2015; Hutt et al., 2018, 2016).

#### Linear stability analysis of the mean-field equations

Fixed points 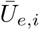 of the mean-field equations satisfy

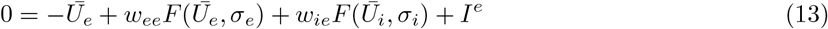

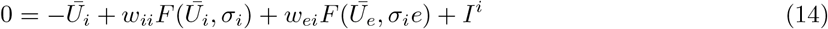

Linearizing about the steady state values of 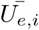 yields the system

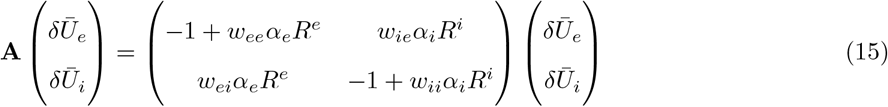

with 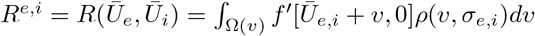. The system’s stability is given by the eigenvalues of the Jacobian **A**. Define

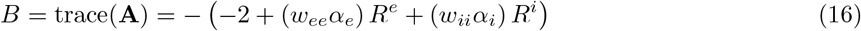

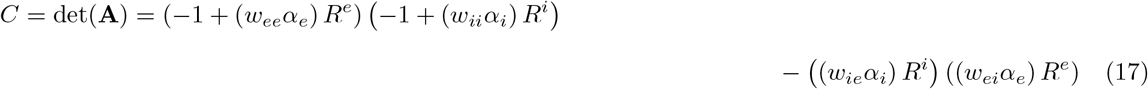

Eigenvalues of **A** are thus given by

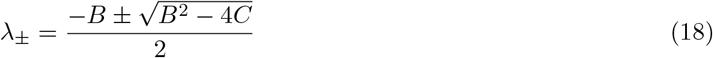

#### Bifurcation analysis with varying excitatory input

We investigate bifurcation properties as a function of *I*^*e*^. In Supplementary Figure S3**(a)**, multi-stability, as denoted by the bold border, is determined by testing for the presence of multiple fixed points at *I*^*e*^ ranging from -15.625:0.625:-6.250, a range encompassing the range for multi-stability shown in Figure 4 (noting *I*^*e*^ = *I*_*e*_ + *I*(*t*)).

## Code Accessibility

The code generating the primary figures is available at https://github.com/Valiantelab/LostNeuralHeterogeneity. Additional code used is available upon request to the authors.

## Supplementary Figures

**Supplementary Figure S1.**
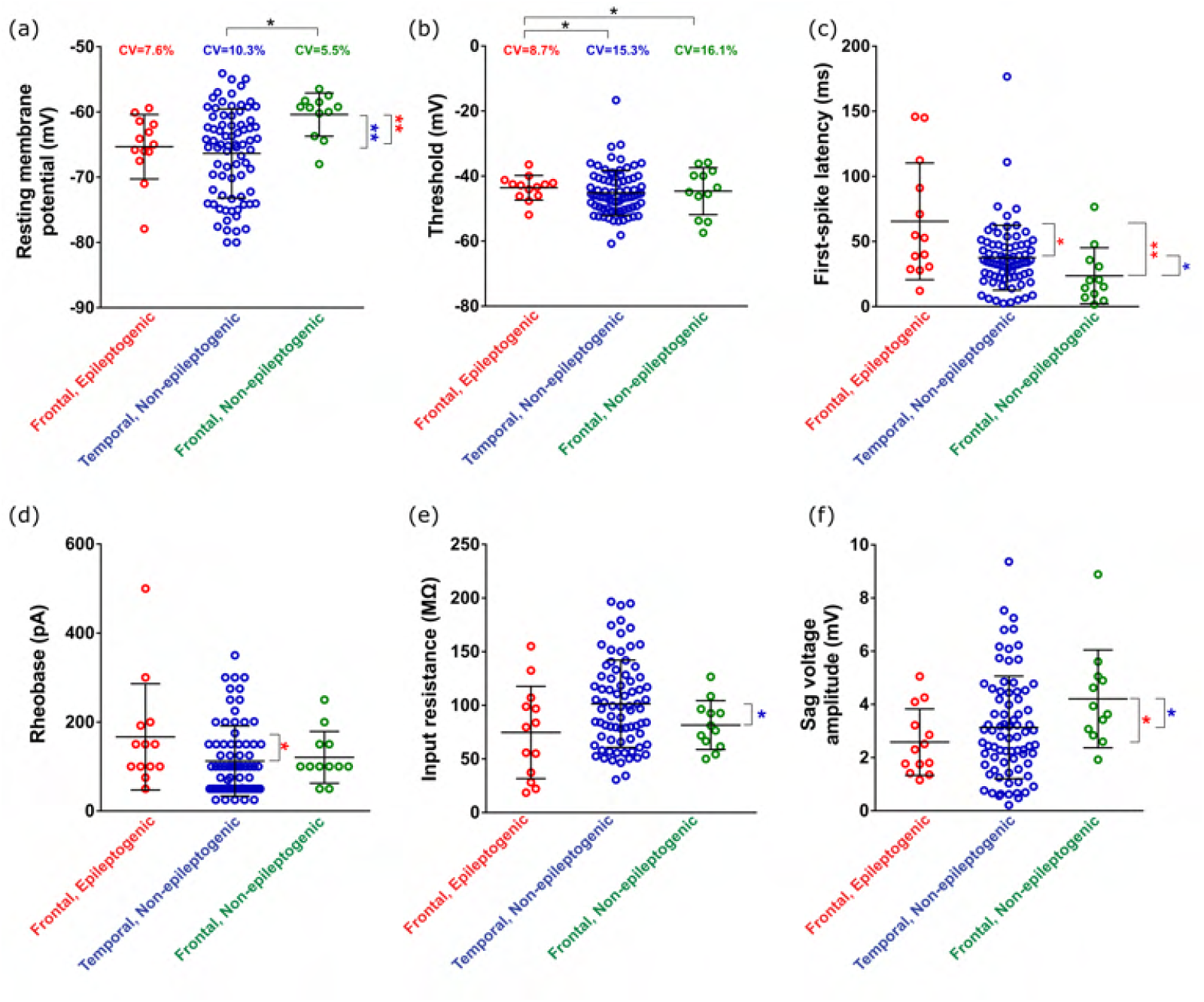
Details from electrophysiological recordings. **(a)**: Distribution of resting membrane potentials (RMP) in our three neuronal populations. Mean RMP is significantly increased in the frontal, non-epileptogenic (n=12) population compared to both the frontal, epileptogenic (n=13; p=0.003; Non-parameteric Mann-Whitney test) and temporal, non-epileptogenic (n=77; p=0.002) populations. Coefficient of variation (CV) of these populations is significantly increased in the temporal, non-epileptogenic population compared to the frontal, non-epileptogenic population (p=0.03; two sample coefficient of variation test). **(b)**: Distribution of threshold voltages in our three neuronal populations. No significant differences between mean threshold voltages were observed (unpaired t test with Welch’s correction). The CV of the threshold voltage in the frontal, epileptogenic population was significantly lower than in the temporal, non-epileptogenic population (p=0.04) and than in the frontal, non-epileptogenic population (p=0.04). **(c)** Distribution of first-spike latencies (time between stimulus application and first spike) in our three neuronal populations. Mean latency is significantly lower in the temporal, non-epileptogenic population compared to the frontal, epileptogenic population (p=0.03; Non-parametric Mann-Whitney test), and mean latency is significantly lower in the frontal, non-epileptogenic population compared to both the frontal, epileptogenic (p=0.0045) and temporal, non-epileptogenic (p=0.02) populations. **(d)** Distribution of rheobases (minimal input current required to elicit first spike) in our three neuronal populations. The mean rheobase of the temporal, non-epileptogenic population is significantly lower compared to the frontal, epileptogenic population (p=0.045; Non-parametric Mann-Whitney test). **(e)** Distribution of input resistances in our three neuronal populations. Mean input resistance of the frontal, non-epileptogenic population is significantly lower compared to the temporal, non-epileptogenic population (p=0.02; Unpaired t-test with Welch’s correction). **(f)** Distributions of sag voltage amplitudes in our three neuronal populations. Mean sag voltage is significantly increased in the frontal, non-epileptogenic population compared to both the frontal, epileptogenic (p=0.01, Non-parametric Mann-Whitney test) and temporal, non-epileptogenic (p=0.04) populations. In all panels, plotted bars indicate mean *±* SD.

**Supplementary Figure S2.**
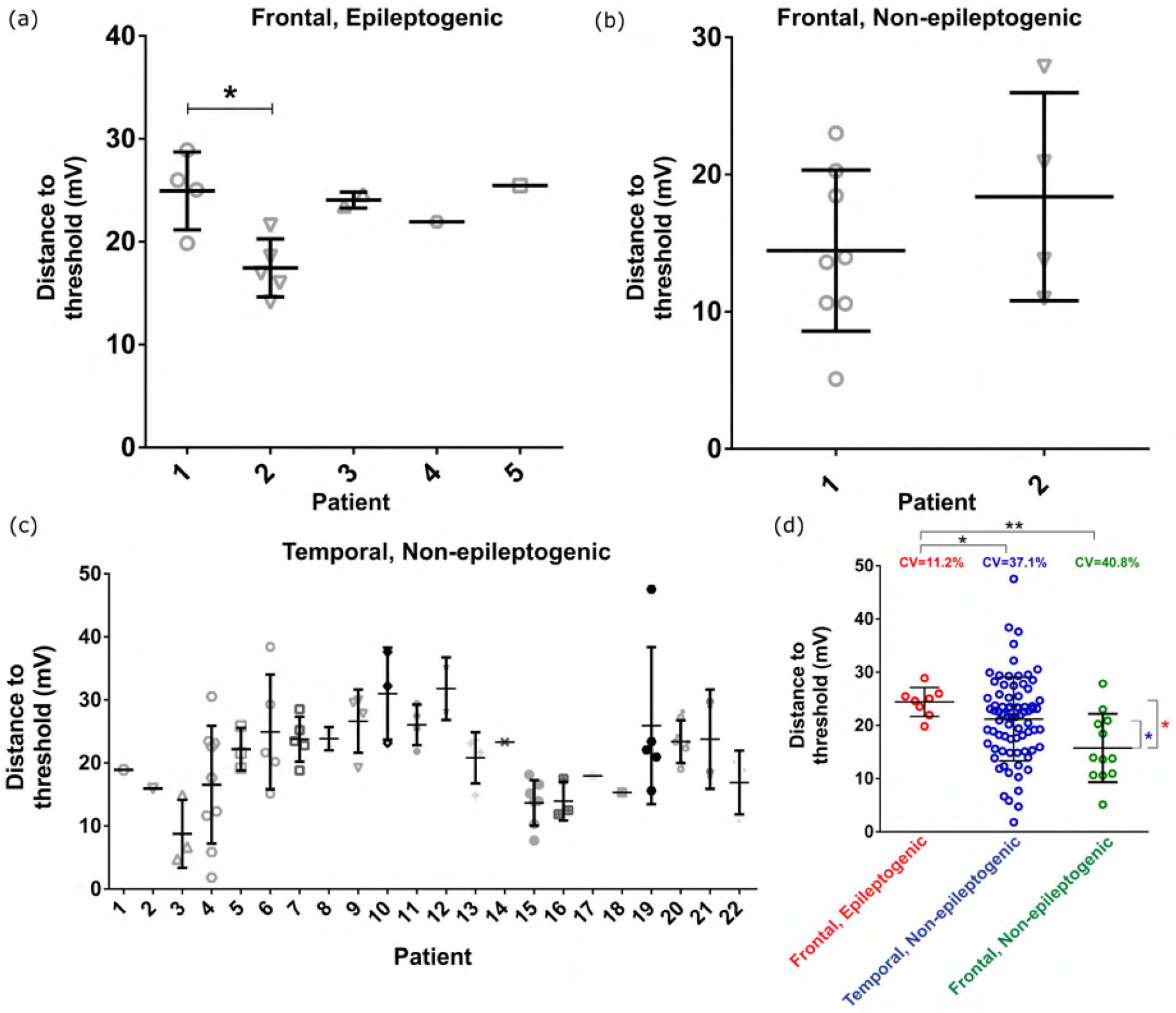
Distance to threshold (DTT) heterogeneities are not skewed by patient-to-patient variability. **(a)**: Frontal, epileptogenic data split by patient. The DTT is significantly different only between Patients 1 and 2 (p=0.04, Dunn’s multiple comparison test; following p=0.02, Kruskal-Wallis test; note: the Kruskal-Wallis test is only performed on patients with *>* 2 cells). This difference would be expected to *increase* the heterogeneity in the frontal, epileptogenic population; nonetheless, its CV is still significantly lower than our two non-epileptogenic populations (see Figure 1**(a)**). **(b)**: Frontal, non-epileptogenic data split by patient. The DTTs are not significantly different (p*>*0.05, non-parametric Mann Whitney test). **(c)**: Temporal, non-epileptogenic data split by patient. No patient’s DTT is statistically different (p*>*0.05, Kruskal-Wallis test; note: the Kruskal-Wallis test is only performed on patients with *>* 2 cells). **(d)**: To confirm our intuition regarding the results presented in panel **(a)** is correct, we replicate the analysis of Figure 1**(a)** but after removing Patient 2’s data from the frontal, epileptogenic population. As expected, the CV of the altered population is decreased, and continues to remain significantly lower than the CVs of the temporal (p=0.02, two-sampled coefficient of variance test) and frontal (p*<*0.01) populations. These results precisely align with recent multi-patch data from human cortex (Planert et al., 2021) highlighting that between-individual variability in biophysical properties was smaller than within-individual variability. Therefore the CVs we measure here are more related to within-individual heterogeneity than between-individual heterogeneity, and thus increased CV is in fact more representative of increased intrinsic biophysical heterogeneity (Planert et al., 2021; Moradi Chameh et al., 2021), than between-subject variability.

**Supplementary Figure S3.**
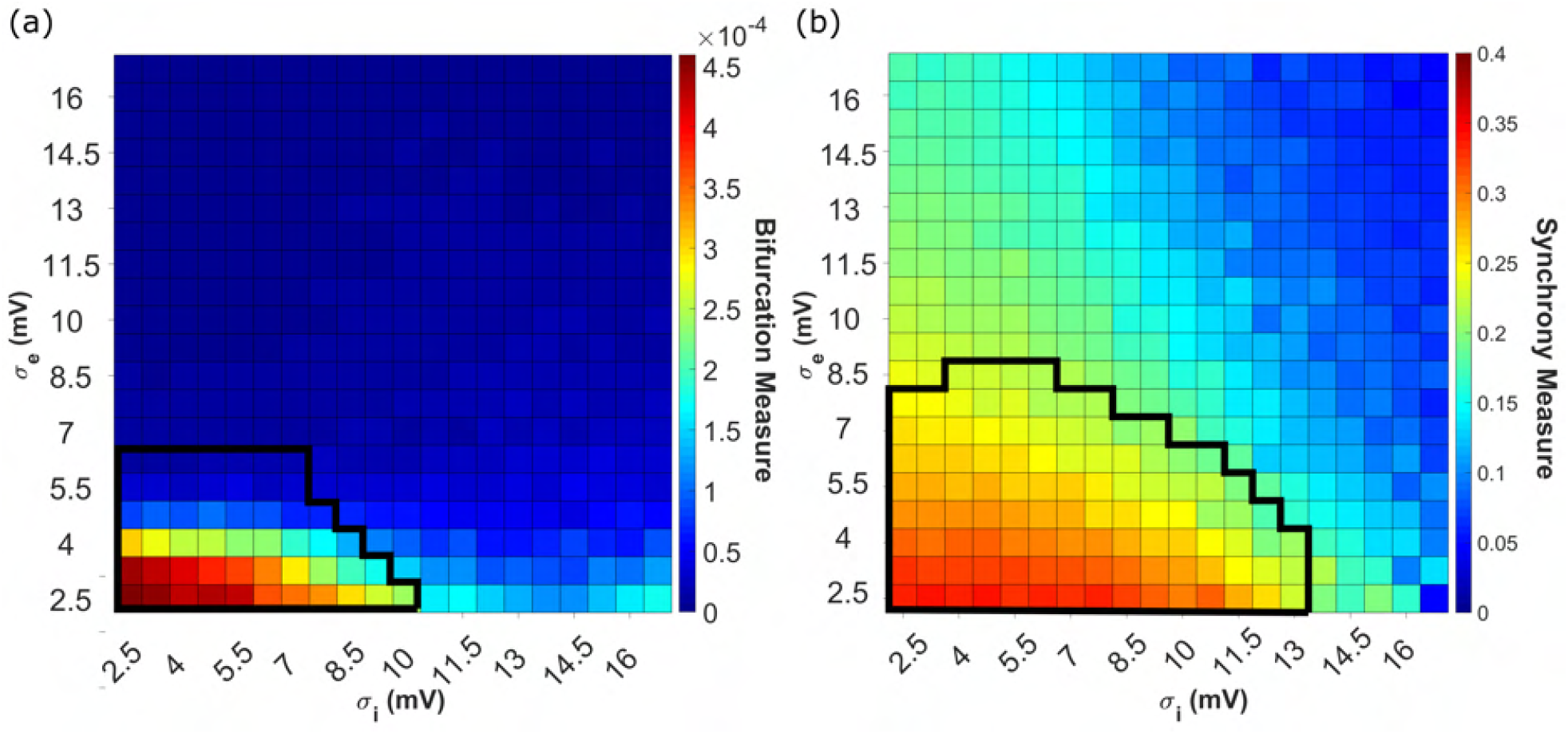
Exploration of a larger *σ*_*e*_ and *σ*_*i*_ parameter space highlights the asymmetric effects of excitatory and inhibitory heterogeneity on sudden transitions into synchrony. **(a)**: Visualization quantifying the tendency for spiking networks to undergo a sudden and notable increase in excitatory synchrony over time, when subjected to a linearly increasing input as in Figures 2, 3, and 4 (but over 2048 as opposed to 2500 ms), via the Bifurcation Measure *B*. Results are shown averaged over 10 independent simulations. Bolded region demarcates networks whose mean-field analogues exhibit any multi-stability from those that do not (remainder of heatmap). **(b)**: Dynamics of spiking networks with a constant external input (*I*(*t*) = 15.625) where either synchronous or asynchronous activity can arise. The excitatory synchrony is quantified via the Synchrony Measure taken over the final 1000 ms of a 2048 ms simulation, and the presented value is averaged over five independent simulations. The bolded region demarcates networks whose mean-field analogues have an unstable oscillator from those that have a stable oscillator (remainder of heatmap) as their lone fixed point when *I*(*t*) = 15.625.

**Supplementary Figure S4.**
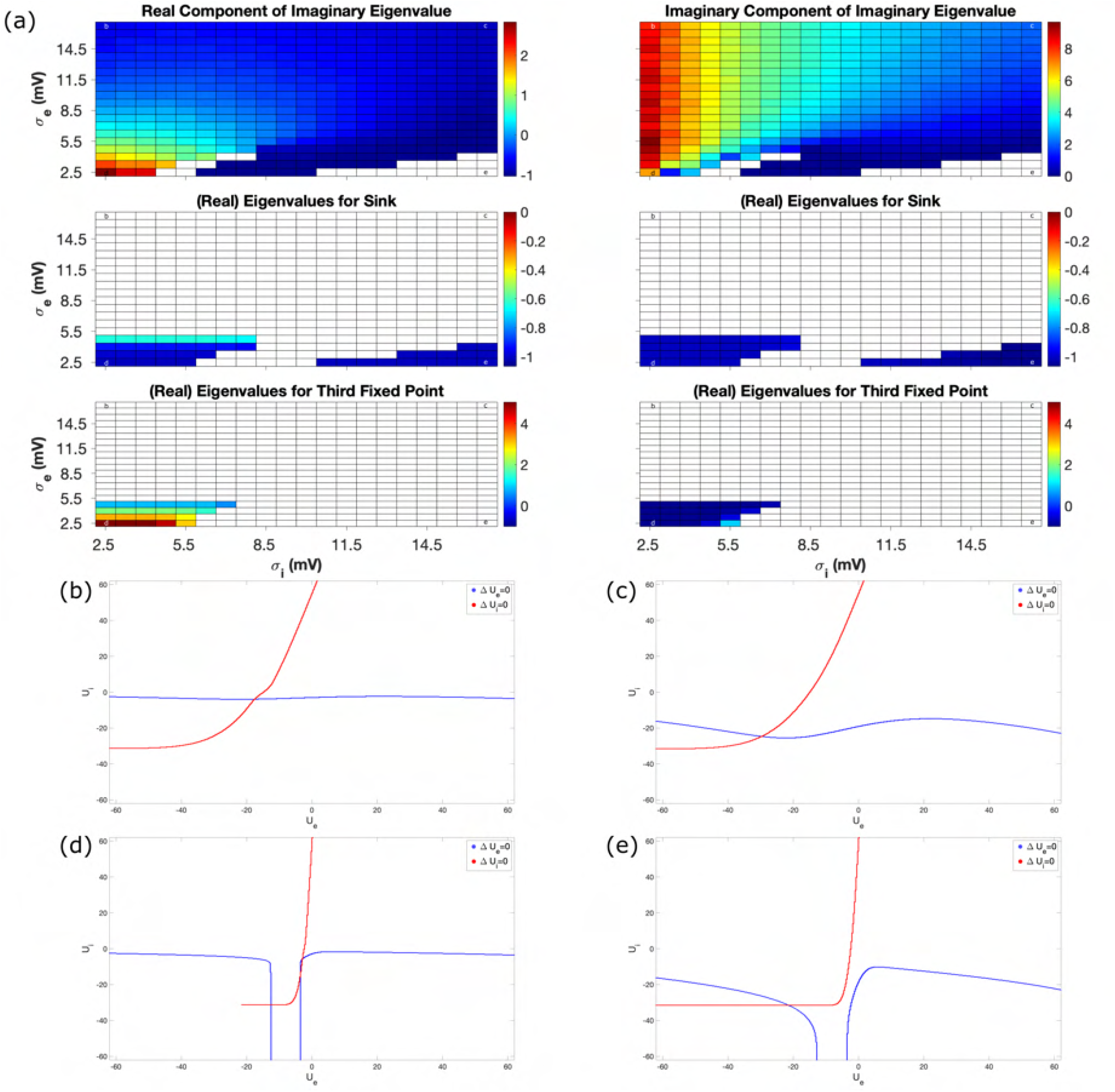
Fixed points and eigenvalues of mean-field equations for *I*(*t*) = 3.125. **(a)**: The mean-field system with this *I*(*t*) value can yield multiple fixed points: we calculate their eigenvalues, sort them by their classifications, and visualize these eigenvalues via heatmaps. In this example, we see that multiple fixed points arise only when both *σ*_*e*_ and *σ*_*i*_ are low (i.e. the bottom-left of the heatmap). **(b-e)**: Fixed points are determined by finding the intersections of the *U*_*e*_ and *U*_*i*_ nullclines, visualized for the corners of our heatmap (top-left in panel **(b)**, top-right in panel **(c)**, bottom-left in panel **(d)**, and bottom-right in panel **(e)**). Multiple fixed points correspond with multiple intersections of these curves, as seen exclusively in panel **(d)**.

**Supplementary Figure S5.**
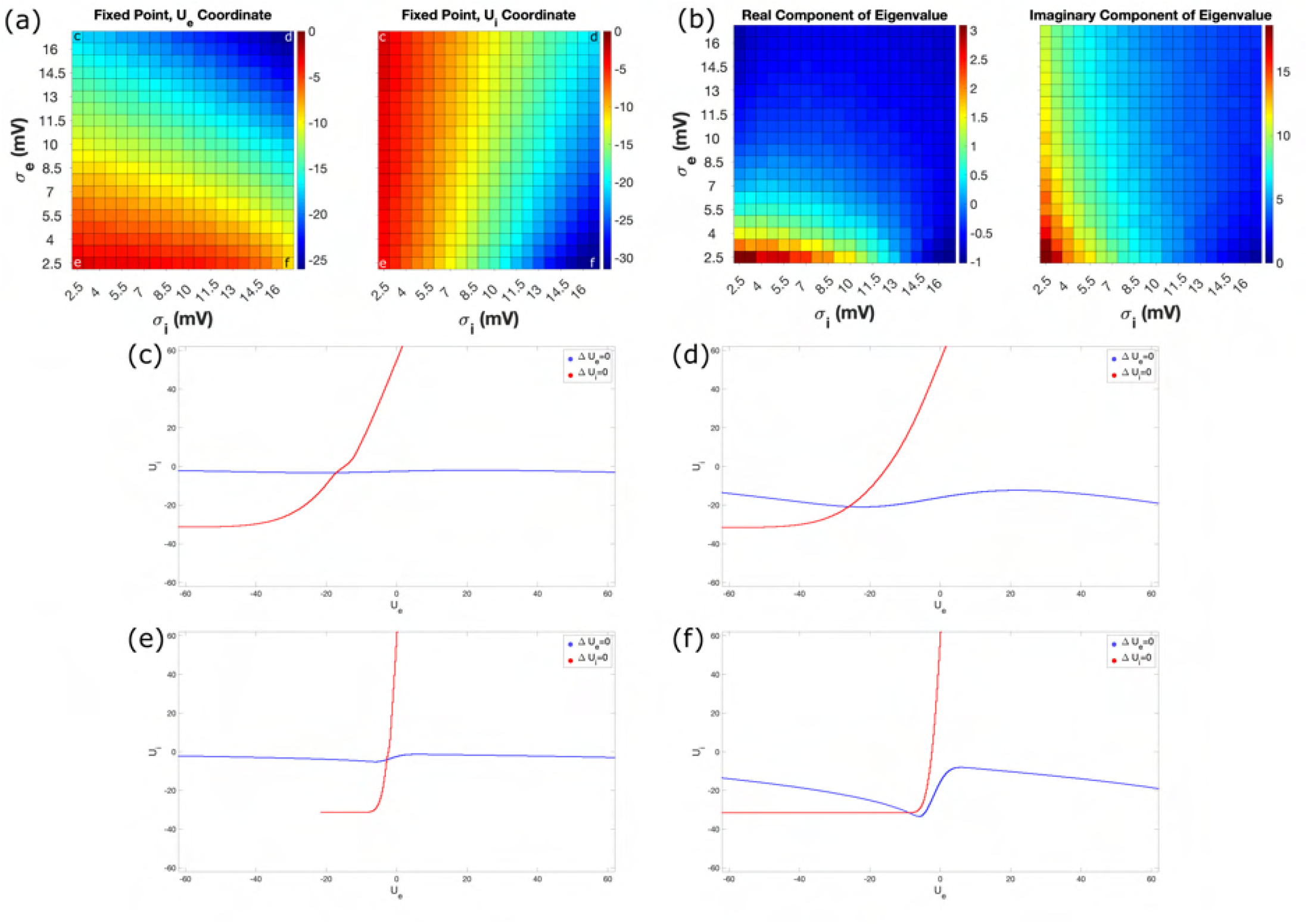
Fixed points and eigenvalues of mean-field equations for *I*(*t*) = 15.625. **(a)**: As all mean-field systems in our parameter space yield a single fixed point when *I*(*t*) = 15.625, we visualize the *U*_*e*_ and *U*_*i*_ coordinates of this fixed point using a heatmap. **(b)**: Each fixed point has imaginary eigenvalues, which we visualize by plotting the real and imaginary components of the eigenvalue associated with the fixed point in a heatmap. **(c-f)**: Fixed points are determined by finding the intersections of the *U*_*e*_ and *U*_*i*_ nullclines, visualized for the corners of our heatmap (top-left in panel **(c)**, top-right in panel **(d)**, bottom-left in panel **(e)**, and bottom-right in panel **(f)**).

**Supplementary Figure S6.**
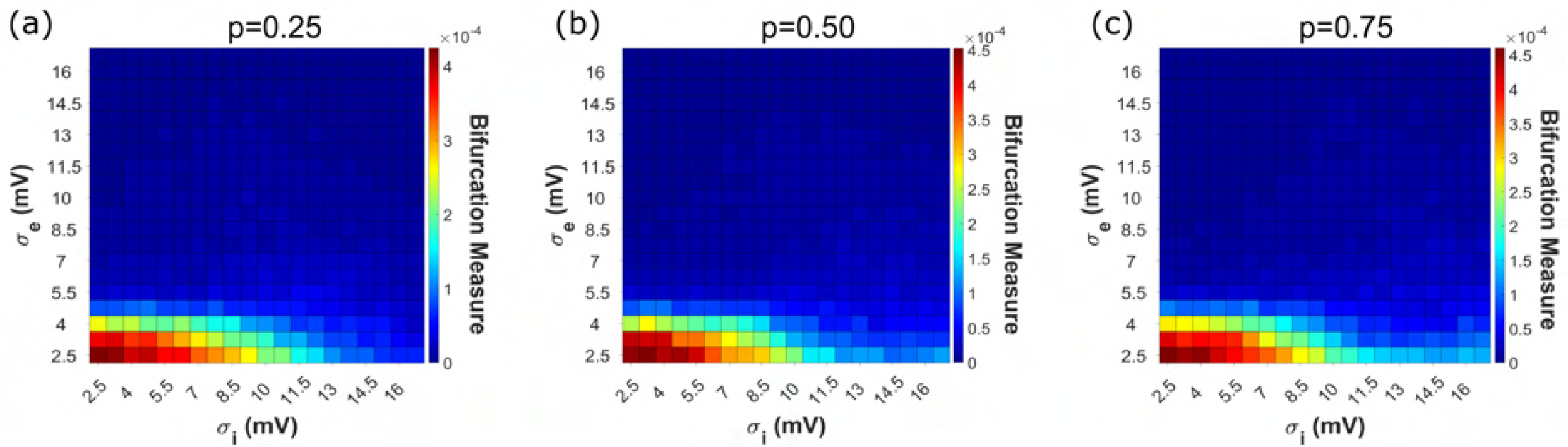
Dynamics of spiking networks are robust to more sparse connectivity paradigms. Bifurcation Measure *B* pattern over our parameter space remains similar with *p* = 0.25 (panel **(a)**), *p* = 0.50 (panel **(b)**), and *p* = 0.75 (panel **(c)**), when compared to the case of *p* = 1.00 seen in Figure S3**(a)**. In each case the “asymmetry” in the effects of *σ*_*e*_ and *σ*_*i*_ is preserved. Heatmaps present results averaged over ten independent simulations.

**Supplementary Figure S7.**
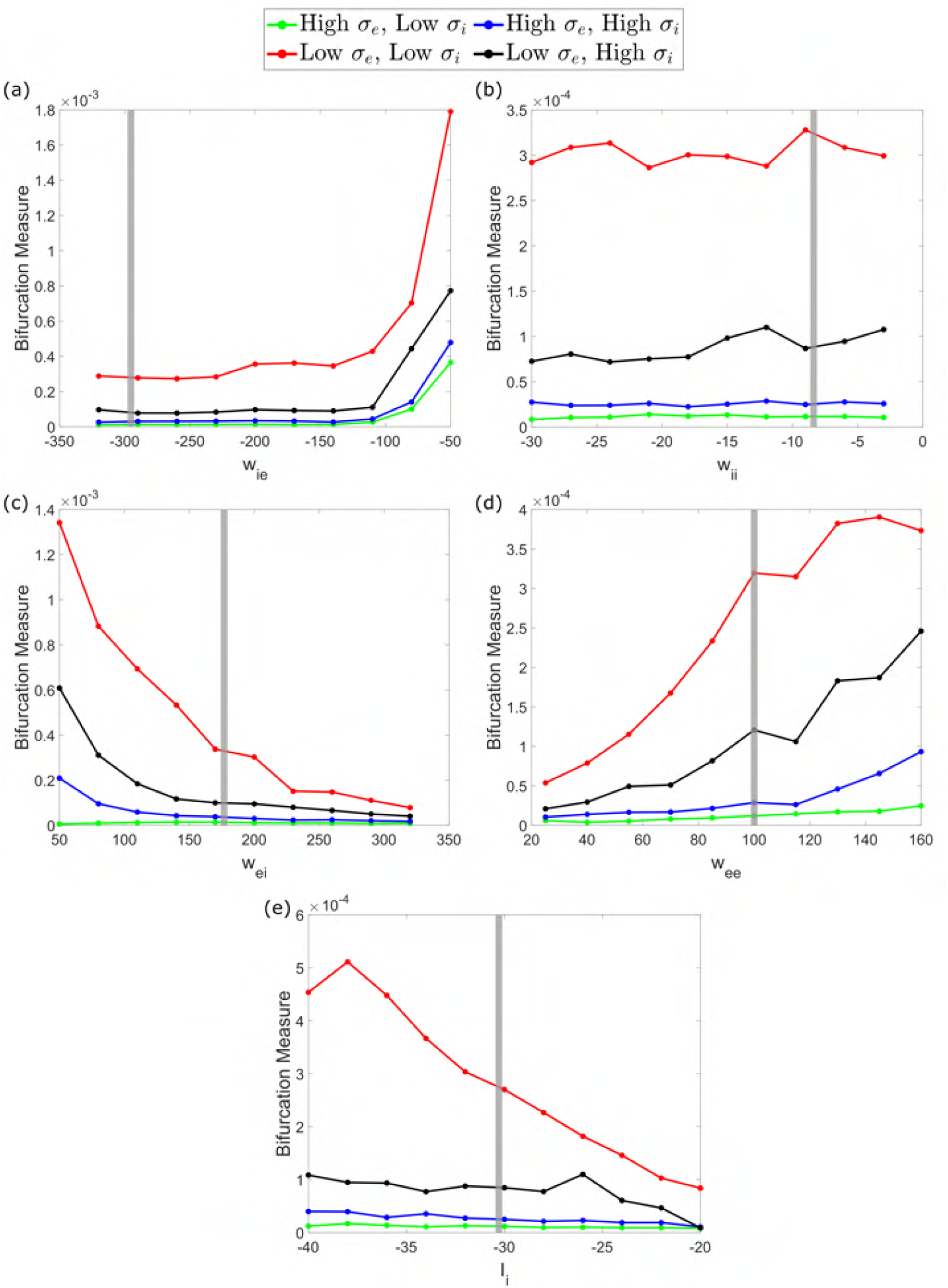
Network dynamics are robust to a range of parameters. **(a-d)**: Examination of changes to network dynamics, as quantified via the Bifurcation Measure, caused by varying a single synaptic weight (*w*_*ie*_ in **(a)**, *w*_*ii*_ in **(b)**, *w*_*ei*_ in **(c)**, and *w*_*ee*_ in **(d)**) or the baseline inhibitory drive (*I*_*i*_ in **(e)**). Vertical grey bar represents the default value as given in Table 1. Values of high/low heterogeneity correspond with those used in exemplar networks in Figures 3 and 4. The Bifurcation Measure is always highest when both *σ*_*e*_ and *σ*_*i*_ are low (red trace), and the other traces (each representing a scenario where at least one of *σ*_*e*_ or *σ*_*i*_ is high) rarely exceed the default Bifurcation Measure of the low *σ*_*e*_ and *σ*_*i*_ case (approximately 3×10^*−*4^). This indicates “sudden transitions” into synchronous dynamics on the magnitude of that seen in Figure 3**(c)** and Figure 4**(c)** occur preferentially in the case of both low *σ*_*e*_ and *σ*_*i*_, even for variations of these parameters. The preserved relationship between the four scenarios represented by the different traces (low *σ*_*e*_ and low *σ*_*i*_ always yielding the highest bifurcation measure, followed by low *σ*_*e*_ and high *σ*_*i*_, followed then by very similar values in both high *σ*_*e*_ scenarios) is further evidence of the robustness of the patterns observed in the results of Figures 3 and 4. Each data point represents an average over 10 independent simulations.

**Supplementary Figure S8.**
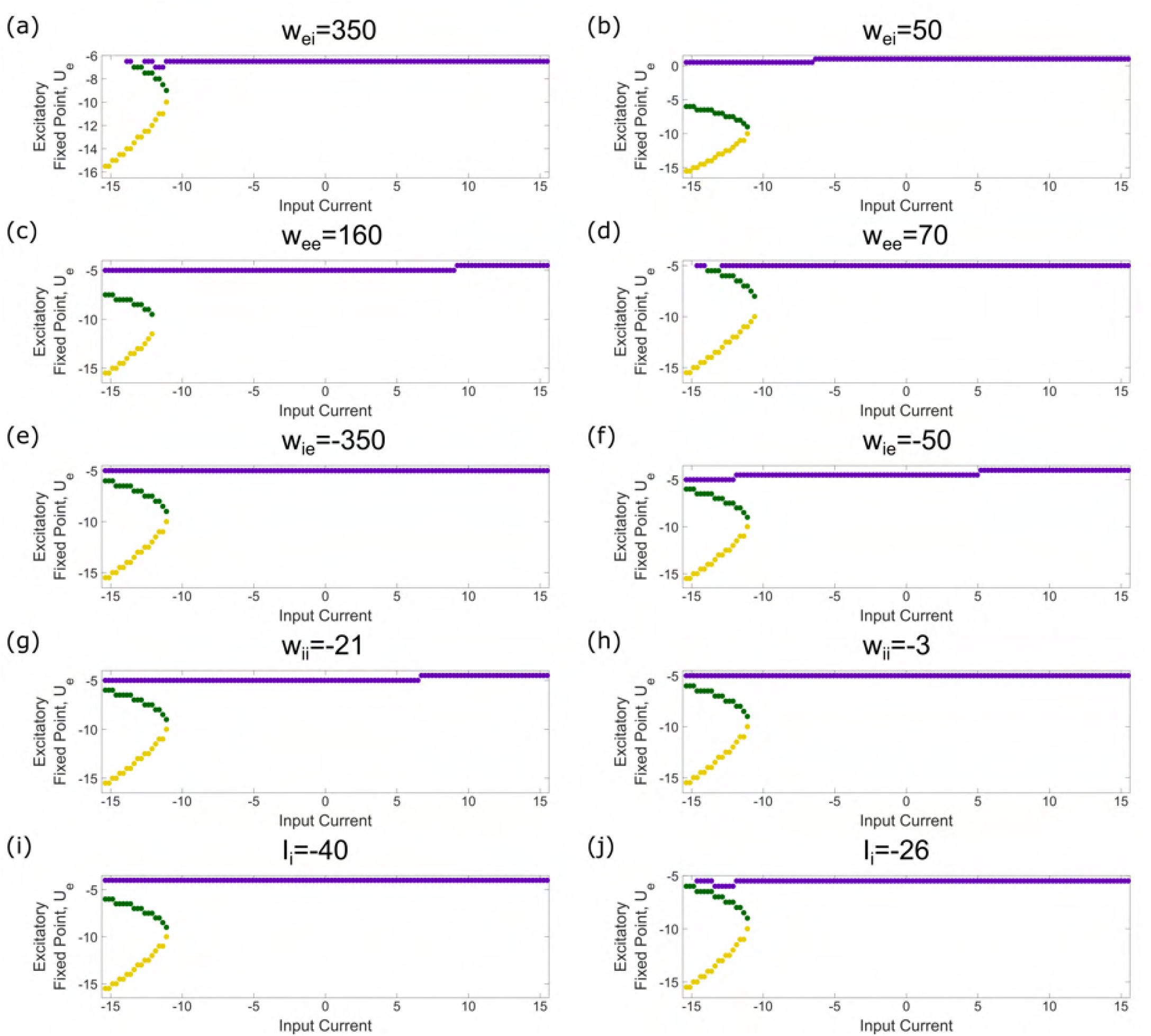
Bifurcation structures are robust to a range of parameters. To confirm the robustness of our spiking network dynamics implies similar robustness in our mean-field systems, we performed bifurcation analyses similar to those in Figure 4 (but with less numerical precision due to computational constraints). Similarly to Supplementary Figure S7 we varied the parameters individually, and showcase examples at high and low extremes for each parameter that clearly preserve the unique bifurcation structure seen in Figure 4**(c)**. This is done for *w*_*ei*_ in **(a-b)**, *w*_*ee*_ in **(c-d)**, *w*_*ie*_ in **(e-f)**, *w*_*ii*_ in **(g-h)**, and *I*_*i*_ in **(i-j)**.

